# RAB18 Orchestrates Signaling Protein Ciliary Homeostasis by Facilitating BBSome Diffusion through the Transition Zone for Ciliary Entry

**DOI:** 10.1101/2025.02.04.636545

**Authors:** Wei-Yue Sun, Jingchao Zhang, Yan Wei, Linda S. Zhao, Zhen-Chuan Fan

## Abstract

The BBSome acts as an intraflagellar transport (IFT) cargo adaptor that connects transmembrane and membrane-anchored signaling proteins to the process of IFT within cilia. This function is critical for ciliary homeostasis maintenance of signaling proteins, allowing for successfully sensing and transducing extracellular stimuli inside the cell. It is established that, upon reaching the proximal ciliary region just above the transition zone (TZ), a diffusion barrier, via retrograde IFT, the BBSome, carrying signaling proteins, disengages from IFT and diffuses through the TZ for ciliary retrieval. This TZ passage event is promoted by the Arf-like 3 GTPase, which utilizes the BBSome autonomously of IFT association as its major effector. While it is known that the intact BBSome is recruited to the basal bodies before entering cilia, the precise mechanism by which it crosses the TZ to access cilia remains elusive. In our study on *Chlamydomonas reinhardtii*, we disclose that the Ras-associated binding (Rab) GTPase RAB18 is concentrated in a basal body region that overlaps with the BBSome. Positioned at this location, RAB18 in a GTP-bound state (RAB18-GTP) is anchored to the membrane, allowing it to diffuse into cilia. Throughout this process, the BBSome functions as a RAB18 effector, binding to RAB18-GTP and subsequently being recruited to diffuse through the TZ for ciliary entry in an IFT-independent manner. This collaboration between the BBSome and RAB18 is pivotal in maintaining the signaling protein phospholipase D ciliary homeostasis, providing a precise mechanistic understanding of the inward BBSome TZ passage essential for proper ciliary signaling.

**Significance statement:** Although certain ciliary signaling proteins cross the transition zone (TZ) for ciliary removal via the RABL2-ARL3 module-mediated outward BBSome TZ diffusion pathway, how the BBSome passes the TZ for ciliary entry remains unclear. Here, we show that RAB18, in its GTP-bound state, anchors to the membrane at the basal bodies. As a RAB18 effector, the BBSome is recruited to cross the TZ for ciliary entry via lateral transport between plasma and ciliary membranes. Once inside cilia, the BBSome integrates into anterograde IFT trains and subsequently functions as a phospholipase D (PLD) adaptor, ensuring PLD removal from cilia. This study highlights RAB18’s pivotal role in facilitating BBSome entry into cilia, unveiling a regulatory mechanism crucial for maintaining ciliary signaling protein homeostasis.

## Introduction

The term cilium and the flagellum are interchangeable, referring to the axonemal microtube-based subcellular organelle that protrudes from the surface of eukaryotic cells. Cilia work as sensory antennas, detecting and transducing the extracellular stimuli into the cell and playing an indispensable role in maintaining various physiological and developmental signaling pathways (1, 2). To fulfil these roles, ciliary transmembrane G protein-coupled receptors (GPCRs), ion channels, and their membrane-anchored downstream signaling components uphold ciliary dynamics by trafficking along the axoneme powered by motor protein-driven intraflagellar transport (IFT) trains, which consists of repeating units of IFT-A (6 subunits) and -B (16 subunits subdivided into IFT-B1 and -B2 entities) complexes (3–6). During IFT, the BBSome, a multi-protein complex of several Bardet-Biedl syndrome (BBS) proteins, acts as an IFT cargo adaptor, bridging signaling proteins to IFT trains (7–10). Defects in BBSome assembly, composition, or ciliary cycling disrupts the proper ciliary localization of signaling proteins, leading to their abnormal loss or accumulation in cilia (7, 8, 10–13). Such disruptions ultimately result in ciliopathic BBS (14, 15).

Across ciliated species, small GTPases serve as molecular switches, mediating various stages of BBSome ciliary cycling (9, 16–23). In *Chlamydomonas reinhardtii,* the Rab-like 5 (RABL5) GTPase IFT22 binds the ARF-like 6 (ARL6) GTPase BBS3 to form the heterodimeric BBS3/IFT22 (16). This complex binds the BBSome via direct interaction with BBS3, recruiting it to the basal bodies when both BBS3 and IFT22 are in their GTP-bound state (16). Upon reaching the ciliary tip via anterograde IFT, the BBSome detaches from IFT trains and undergoes remodeling in the ciliary tip matrix via a disassembly-reassembly process (20, 24). The remodeled BBSome then acts as a effector of ARL13, anchoring to the ciliary tip membrane to facilitate ciliary phospholipase D (PLD) binding through the BBS3-ARL13 module (18, 22). Following ARL13 hydrolysis, the PLD-laden BBSome loads onto retrograde IFT trains and completes a U-turn at the ciliary tip (22). Upon reaching at the proximal ciliary region right above the ciliary diffusion barrier, the transition zone (TZ), via retrograde IFT, the PLD-laden BBSome dissociates from retrograde IFT trains and diffuses through the TZ for ciliary retrieval as an effector of ARL3 via the RABL2-ARL3 module (21, 23, 25). In coordination with the RABL2-ARL3 module, the BBSome promotes the crossing of PLD through the TZ for ciliary retrieval, suggesting that lateral transport between ciliary and plasma membranes is the primary mechanism for the BBSome’s movement across the TZ (21, 23, 25). Despite these insights, the exact molecular mechanism underlying the BBSome’s movement across the TZ for ciliary entry remains unclear.

Ras-associated binding (RAB) GTPase RAB18 localizes to the Golgi apparatus, endoplasmic reticulum (ER), lipid droplets, and cytosol of various cell types (26–28). It plays a key role in regulating intercellular membrane trafficking, particularly between the ER and Golgi apparatus (26). This study reveals that *Chlamydomonas* RAB18 is enriched at the basal bodies and anchored to the membrane in its GTP-bound state (RAB18-GDP) to facilitate its diffusion into cilia. During this process, the BBSome acts as a RAB18 effector, binding to RAB18-GTP and being subsequently recruited to diffuse through the TZ for ciliary entry. Once inside cilia, the BBSome integrates into anterograde IFT trains, a process that appears to occur in the proximal ciliary base. This integration allows the BBSome to maintain PLD ciliary homeostasis by acting as an IFT cargo adaptor. These findings bridge a critical gap regarding how the BBSome crosses the TZ for ciliary entry and suggests that RAB18 deficiency may contribute to BBS phenotypes in humans.

## Results

### RAB18 enters cilia through diffusion and does not influence IFT

*Chlamydomonas* RAB18 shares over 49% identities with its orthologues in other ciliated species (*SI Appendix,* Fig. S1*A* and *B*). Unlike several other RAB18 that are phylogenetically closely related in ciliated species, *Chlamydomonas* RAB18 is less closely related to human RAB18 (*SI Appendix,* Fig. S1*C*). Proteomics analysis of *Chlamydomonas* cilia identifies RAB18 as a putative ciliary regulator with an unknown function (29). To investigate the role of RAB18 in IFT and ciliation in *C. reinhardtii*, we attempted to generate a RAB18-knockout mutant in the CC-5325 strain using CRISPR/Cas9 but were unsuccessful (30, 31). We then opted to use the CLiP mutant (LMJ.RY0402.132876, referred to as *rab18-876*), which contains an insertion at the 3’-proximal region of the fifth exon of the *rab18* gene, but found that it is not a true RAB18-null mutant (*SI Appendix,* Fig. S2*A-*2*D*). As an alternative, we investigated the role of *Chlamydomonas* RAB18 in IFT and ciliation by using a knockdown (KD) approach with a highly efficient artificial miRNA system (32). Using the newly developed RAB18 antibody (*SI Appendix,* Fig. S2*A*), we identified a RAB18 KD strain named *RAB18^miR^* in the CC-5325 background, which exhibited a 93% reduction in RAB18 levels compared to parental CC-5325 cells, as confirmed by immunoblotting (Fig. 1*A*). Correspondingly, the RAB18 levels in *RAB18^miR^*cilia showed a proportional reduction (Fig. 1*A*). Notably, *RAB18^miR^*cells displayed normal cilia growth, effectively ruling out a direct role for RAB18 in ciliation (*SI Appendix,* Fig. S3*A* and *B*). Supporting this conclusion, *RAB18^miR^* cells maintained endogenous levels of IFT-A subunits (IFT43 and IFT139), IFT-B1 subunits (IFT22 and IFT70), and IFT-B2 subunits (IFT38 and IFT57) both in whole cell samples and cilia under steady-state conditions (Fig. 1*A*). To determine if RAB18 influences IFT ciliary dynamics, we generated transgenic strains *RAB18^miR^; IFT43:HA:YFP-TG*, *RAB18^miR^; IFT22:HA:YFP-TG*, and *RAB18^miR^; IFT38:YFP-TG*, which expressed IFT43, IFT22, or IFT38 fused to hemagglutinin (HA) and/or yellow fluorescent protein (YFP) at their C-termini (IFT43-HA-YFP, IFT22-HA-YFP, and IFT38-YFP) in the *RAB18^miR^* strain. These tagged proteins were expressed at comparable levels in the parental CC-5325 strain, resulting in control strains *RAB18; IFT43:HA:YFP-TG*, *RAB18; IFT22:HA:YFP-T*G, and *RAB18; IFT38:YFP-TG* (*SI Appendix,* Fig. S3*C*). In *RAB18^miR^* cells, these proteins entered cilia and dominated over the endogenous proteins (Fig. 1*B*). Notably, the combined abundance of endogenous proteins and their tagged counterparts equaled the endogenous levels observed in CC-5325 cilia, indicating that the RAB18’s ciliary content is tightly controlled in *C. reinhardtii* (Fig. 1*B*). Total internal reflection fluorescence (TIRF) microscopy confirmed that these tagged proteins underwent normal bidirectional IFT, excluding RAB18 from a role in regulating IFT ciliary dynamics (Fig. 1*C* and *D* and Movies S1-S6). To explore how *Chlamydomonas* RAB18 enters cilia, we expressed RAB18-HA-YFP in *RAB18^miR^* cells at levels comparable to endogenous RAB18 level of CC-5325 cells, generating the strain *RAB18^miR^; RAB18:HA:YFP-TG* (*SI Appendix,* Fig. S3*D*). This strain retained RAB18-HA-YFP in cilia at 93% of the endogenous RAB18 level (Fig. 1*E*). TIRF microscopy showed that RAB18-HA-YFP entered cilia through a diffusion rather than via IFT and was distributed uniformly alone the entire cilia (Fig. 1*F* and Movie S7). In summary, RAB18 does not impact IFT in both steady and dynamic states. IT enters cilia through diffusion and is distributed throughout the entire ciliary region.

**Figure 1.**
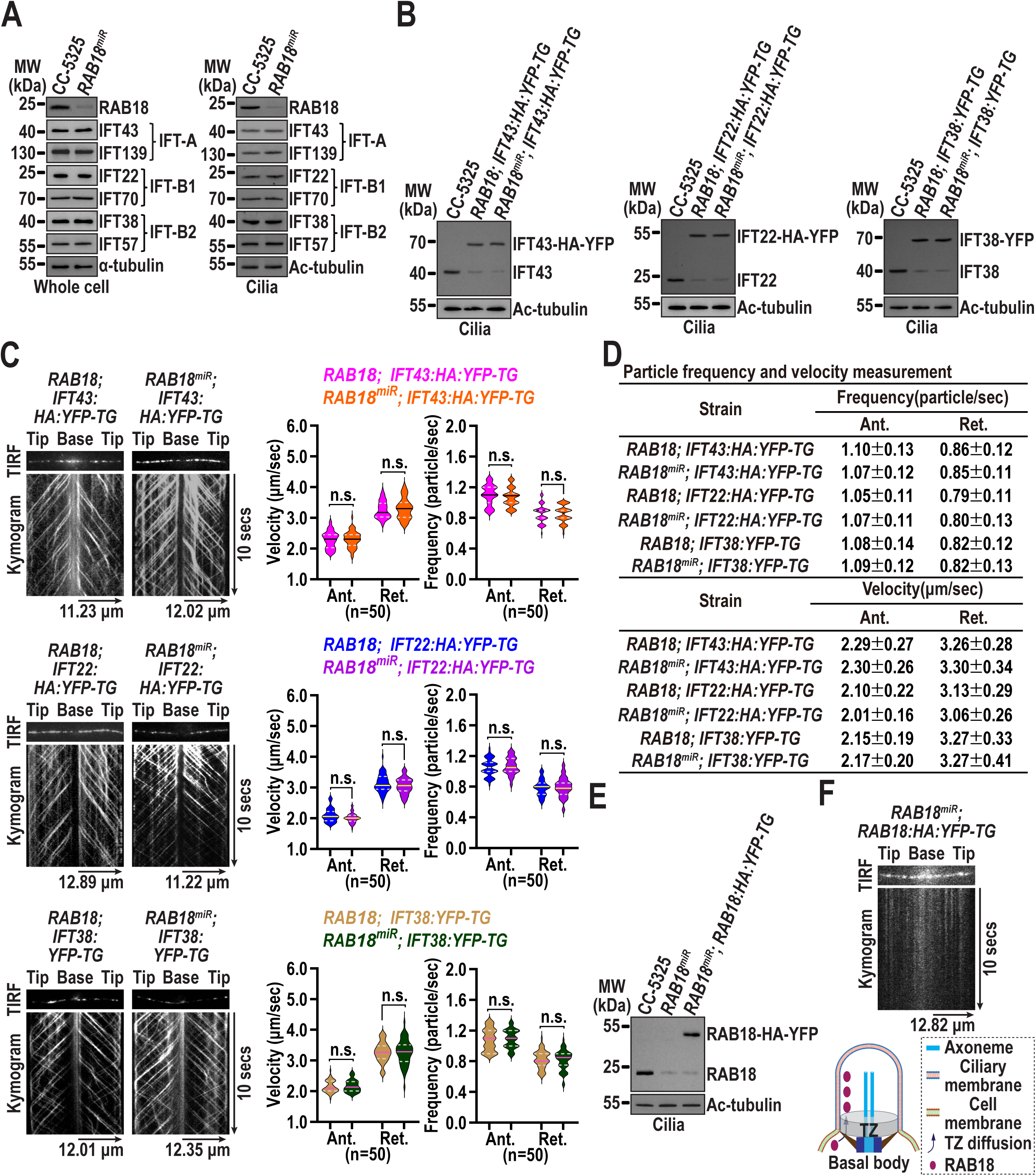
RAB18 enters cilia through diffusion and does not influence IFT. **(A)**. Immunoblots of whole cell samples and cilia from CC-5325 and *RAB18^miR^* cells probed for RAB18, IFT43 and IFT139 (IFT-A), IFT22 and IFT70 (IFT-B1), and IFT38 and IFT57 (IFT-B2). (**B)**. Immunoblots of cilia from three cell groups (as indicated on the top) probed with α-IFT43, α-IFT22, and α-IFT38. **(C)**. Representative TIRF images and corresponding kymograms of the three cell groups shown in panel B (Movies S1-S6, 15 fps). Graphs depicts the velocities and frequencies of YFP- and HA-YFP-labeled proteins trafficking inside cilia. Error bar indicates standard deviation (S.D.) and “n” represents the number of cilia analyzed. Non-significant differences are demoted as n.s. A one sample unpaired student’s *t-*test was used. **(D)**. Numerical values of velocities and frequencies of YFP- and HA-YFP-tagged proteins for ciliary trafficking shown in panel C. **(E)**. Immunoblots of cilia from CC-5325, *RAB18^miR^*, and *RAB18^miR^; RAB18:HA:YFP-TG* cells probed with α-RAB18. **(F)**. Representative TIRF image and corresponding kymogram of the *RAB18^miR^; RAB18:HA:YFP-TG* cell (Movie S7, 15 fps). A schematic representation illustrating how RAB18 crosses the TZ for diffusing into cilia is shown below. For panels **A**, **B**, and **E**, α-tubulin and acetylated (Ac)-tubulin were used to normalize the loading of whole cell samples and cilia, respectively. MW: molecular weight. For panels **C** and **D**, “Ant.” and “Ret.” represent anterograde and retrograde trafficking, respectively. For panels **C** and **F**, time and transport lengths are indicated on the right and on the bottom, respectively. The ciliary base (base) and tip (tip) are labeled.

### RAB18-GTP anchors to the membrane, allowing its diffusion through the TZ and entry into cilia

The GTP-bound state is a crucial prerequisite for RAB GTPases to efficiently anchor to the membrane (33). To examine how nucleotide state influence RAB18 anchoring to the ciliary membrane, we isolated the ciliary membrane, matrix (the content between the ciliary membrane and axoneme), and axonemal fractions from CC-5325 cells in the presence of the nonhydrolyzed GTP analog GTPγS, GDP, or no added nucleotide. GTPγS and GDP lock RAB18 in GTP- and GDP-bound state, respectively. In the presence of GDP or no nucleotide, RAB18 was detected exclusively in the ciliary matrix (Fig. 2*A*). However, with GTPγS, RAB18 localized solely to the ciliary membrane fraction (Fig. 2*A*). These results demonstrate that the GTP-bound state is essential for efficient membrane anchoring of *Chlamydomonas* RAB18. To further confirm this, we expressed RAB18 and its variants carrying the Q67L (GTP-locked, active form) or S22N (GDP-locked/nucleotide-free, inactive form) mutation in bacteria (34–36). Liposome flotation assays using bacterially expressed RAB18 preloaded with GTPγS or GDP revealed that RAB18 bound to liposomes only when preloaded with GTPγS, confirming that membrane anchoring requires the GTP-bound state (Fig. 2*B*). Furthermore, RAB18Q67L exhibited liposome binding, whereas RAB18S22N did not, reinforcing that RAB18Q67L mimics the GTP-bound state that anchors efficiently to membranes (Fig. 2*B*).

**Figure 2.**
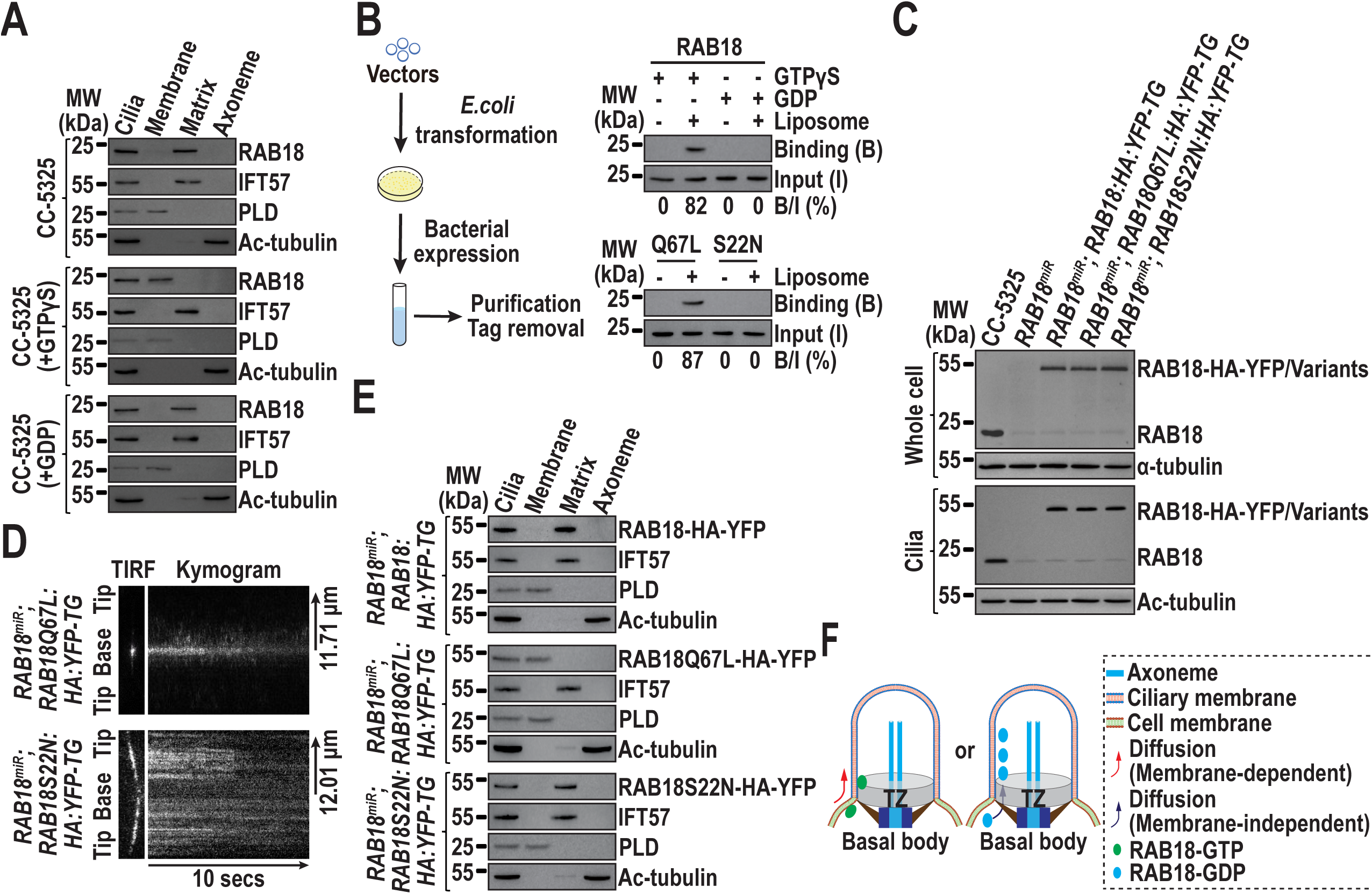
RAB18-GTP is anchored to the membrane for diffusing into cilia. **(A)**. Immunoblots of ciliary fractions from CC-5325 cells in the presence of GTPγS, GDP or none of them probed with α-RAB18, α-IFT57 (ciliary matrix marker), α-PLD (ciliary membrane marker), and Ac-tubulin (axoneme marker). MW: molecular weight. **(B)**. Immunoblots of liposome flotation-captured RAB18 preloaded with GTPγS or GDP (upper) and its variants (bottom) using α-RAB18. RAB18 or its variants along was used as inputs, with the binding/input ratio displayed as a percentile. **(C)**. Immunoblots of whole cell samples and cilia from cells indicated on the top probed with α-RAB18. Alpha-tubulin and Ac-tubulin were used to normalize the loading of whole cell samples and cilia, respectively. MW: molecular weight. **(D)**. Representative TIRF images and corresponding kymograms of cells indicated on the left (Movies S8-S9, 15 fps). Time and transport lengths are indicated on the bottom and on the right, respectively. The ciliary base (base) and tip (tip) are labeled. **(E)**. Immunoblots of ciliary fractions from cells indicated on the left probed with α-YFP, α-IFT57 (ciliary matrix marker), α-PLD (ciliary membrane marker) and Ac-tubulin (axoneme marker). MW: molecular weight. **(F)**. Schematic presentation of how nucleotide state determines RAB18 crosses the TZ for diffusing into cilia in a membrane-dependent (GTP-bound) or a nonmembrane-dependent (GDP-bound) pattern. Only in its GTP-bound form is RAB18 concentrated in the proximal ciliary region upon entry.

To investigate how membrane anchoring influences RAB18 entry into cilia, we expressed the two RAB18 variants fused at their C-terminus to HA-YFP in RAB18 knockdown (*RAB18^miR^*) cells (Fig. 2*C*). This resulted in strains *RAB18^miR^; RAB18Q67L:HA:YFP-TG* and *RAB18^miR^; RAB18S22N:HA:YFP-TG*, which were screened to express the tagged RAB18 variants at the RAB18-HA-YFP level in *RAB18^miR^; RAB18:HA:YFP-TG* cells (Fig. 2*C*). In *RAB18^miR^* cells, RAB18-HA-YFP behaved like the endogenous RAB18 in entering cilia (Fig. 2*C*, also see Fig. 1*E*). Both RAB18Q67L-HA-YFP and RAB18S22N-HA-YFP entered cilia, mimicking the endogenous RAB18 and RAB18-HA-YFP (Fig. 2*C*; also see Fig. 1*E*). TIRF microscopy visualized RAB18Q67L-HA-YFP to diffuse into cilia and primarily concentrated in the proximal ciliary region (Fig. 2*D* and Movie S8). It was entirely localized to the ciliary membrane fraction, as shown by ciliary fractionation assays (Fig. 2*E*). In contrast, RAB18S22N-HA-YFP diffused into cilia, but was distributed along their entire cilia (Fig. 2*E* and Movie S9). Unlike the Q67L mutant, it was exclusively localized to the ciliary matrix fraction (Fig. 2*E*). These findings suggest that RAB18-GTP diffuses into cilia while remaining attached to the membrane (Fig. 2*F*). Conversely, RAB18-GDP likely diffuses into cilia via membrane-independent diffusion across the TZ (Fig. 2*F*). Upon diffusing into cilia, RAB18-HA-YFP resembled the S22N mutant, but not the Q67L mutant, in its distribution along the entire length of cilia (Fig. 1*F*) and its localization within the ciliary matrix (Fig. 2*E*). Given the millimolar GTP concentrations in the cell (37) and micromolar affinity of RAB18 for guanine nucleotides (38), it is likely that RAB18-GTP, rather than RAB18-GDP, diffuses into cilia via lateral transport between the cell and ciliary membranes under physiological conditions. Upon crossing the TZ and reaching the proximal ciliary region, RAB18-GTP undergoes rapid nucleotide exchange, converting to RAB18-GDP. This conversion causes RAB18-GDP to detach from the ciliary membrane and localize within the ciliary matrix.

### RAB18-GTP promotes BBSome passage through the TZ for ciliary entry

In *RAB18^miR^* cells, PLD accumulates in cilia (Fig. 3*A*), predominantly at the ciliary tip as visualized by immunofluorescence staining (Fig. 3*B*). This ciliary tip phenotype was restored to normal, rendering PLD undetectable at the ciliary tip in *RAB18^miR^; RAB18:HA:YFP-TG* cells but not in *RAB18^miR^; RAB18Q67L:HA:YFP-TG* and *RAB18^miR^; RAB18S22N:HA:YFP-TG* cells (Fig. 3*A* and *B*). Our previous study has showed that the ciliary membrane-anchored PLD engages with the BBSome at the ciliary tip via the BBS3-ARL13 module-mediated cargo-BBSome coupling pathway (19, 22). Since RAB18 does not influence IFT (Fig. 1*A-D*), we examined whether RAB18 regulates PLD ciliary dynamics through influencing BBSome abundance in cilia. Compared to parental CC-5325 cells, the BBSome (represented by BBS1, BBS4, BBS5, BBS7, and BBS8) was retained at control levels in whole cell samples from *RAB18^miR^* cells and the strains *RAB18^miR^; RAB18:HA:YFP-TG*, *RAB18^miR^; RAB18Q67L:HA:YFP-TG*, and *RAB18^miR^; RAB18S22N:HA:YFP-TG* (Fig. 3*A*). However, significant reductions in BBSome levels were observed in cilia from *RAB18^miR^* and *RAB18^miR^; RAB18S22N:HA:YFP-TG* cells (Fig. 3*A*). The restoration of normal ciliary BBSome levels by RAB18-HA-YFP, particularly the GTP-locked RAB18Q67L-HA-YFP, identifying RAB18-GTP promotes BBSome entry into cilia (Fig. 3*A*).

**Figure 3.**
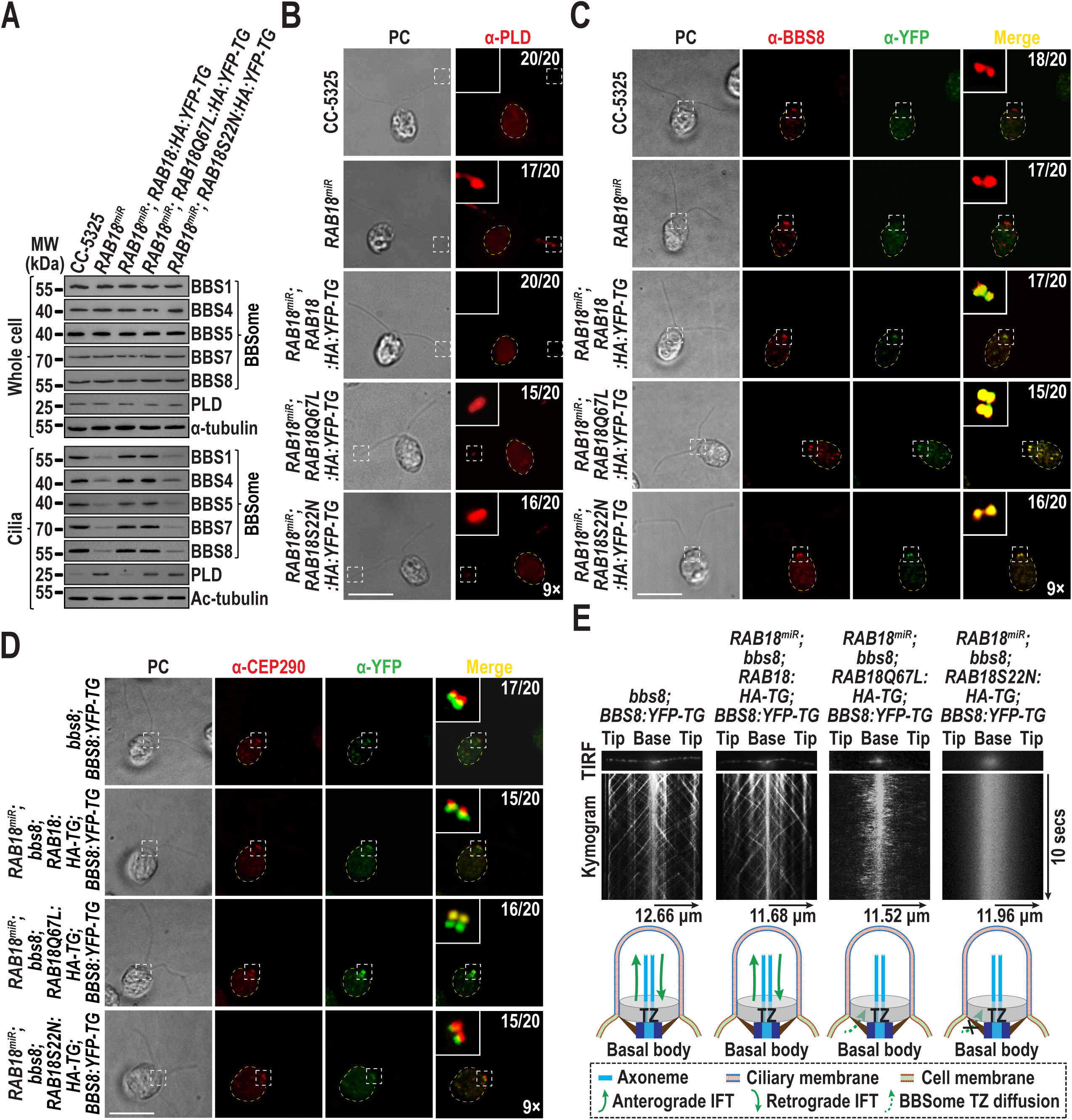
RAB18-GTP promotes BBSome passage through the TZ for entry into cilia. **(A)**. Immunoblots of whole cell samples and cilia from the indicated cells probed for BBSome subunits (BBS1, BBS4, BBS5, BBS7, and BBS8) and PLD. Alpha-tubulin and Ac-tubulin were used to normalize the loading of whole cell samples and cilia, respectively. MW: molecular weight. **(B)**. Immunofluorescence staining of the indicated cells with α-PLD (red). **(C)**. Immunofluorescence staining of the indicated cells with α-BBS8 (red) and α-YFP (green). BBS8 serves as a basal body marker. **(D)**. Immunofluorescence staining of the indicated cells with α-CEP290 (red) and α-YFP (green). CEP290 and BBS8-YFP serve as a TZ and basal body marker, respectively. **(E)**. Representative TIRF images and corresponding kymograms of the indicted cells (Movies S10-S13, 15 fps). A schematic representation of the BBSome undergoing IFT between the ciliary proximal region right above the TZ and the ciliary tip, and the RAB18 nucleotide-dependent movement across the TZ for ciliary entry, is shown below. For panels **B**, **C**, and **D**, phase contrast (PC) images of cells are included. Insets highlight the proximal ciliary region right above the TZ and basal bodies. Inset magnifications (9 times) are displayed. Scale bars: 10 µm. Cell numbers out of 20 cells are listed for representing cells that are signal positive in the basal body region and the ciliary base right above the TZ.

Compared to the parental CC-5325 cells, the RAB18 knockdown strain *RAB18^miR^* has RAB18 levels sufficiently low to be undetectable at the basal bodies by immunofluorescence staining (*SI Appendix,* Fig. S4). The HA-YFP tagged RAB18 and its variants effectively mimic the endogenous RAB18 in targeting the basal bodies, indicating that the tags and the introduced mutations do not interfere with RAB18 basal body distribution and function (*SI Appendix,* Fig. S4). Neither RAB18 knockdown nor its replacement with RAB18Q67L-HA-YFP or RAB18S22N-HA-YFP disrupted BBSome (represented by BBS8) targeting to the basal bodies normally (Fig. 3*C* and *SI Appendix,* Fig. S4). At the basal bodies, RAB18 in either GTP- or GDP-bound state co-localized with the BBSome (Fig. 3*C*). Notably, the BBSome and the Q67L mutant were also confined together, likely co-localizing in the proximal ciliary region (Fig. 3*C*, *SI Appendix,* Fig. S4). To precisely determine the BBSome localization relative to the TZ in the presence of RAB18 variants, we generated strains *RAB18^miR^; RAB18:HA-TG*, *RAB18^miR^; RAB18Q67L:HA-TG*, and *RAB18^miR^; RAB18S22N:HA-TG.* These three strains were crossed with *bbs8; BBS8:YFP-TG*, eventually yielding the strains *RAB18^miR^; bbs8; RAB18:HA-TG; BBS8-YFP-TG*, *RAB18^miR^; bbs8; RAB18Q67L:HA-TG; BBS8-YFP-TG*, and *RAB18^miR^; bbs8; RAB18S22N:HA-TG; BBS8-YFP-TG*) (*SI Appendix,* Fig. S5*A* and *B*) (21). In these strains, BBS8-YFP and RAB18-HA (or its variants) replaced their endogenous counterparts to enable proper immunofluorescence staining (*SI Appendix,* Fig. S5*A* and *B*). The presence of either the Q67L or S22N mutant does not affect BBSome targeting to the basal bodies, as visualized below the CEP290-labeled TZ (Fig. 3*D*). However, it was the Q67L mutant that confines the BBSome in the proximal ciliary region, overlapping with the CEP290-labeled TZ (Fig. 3*D*). As a result, neither the loss of RAB18 nor the presence of its variants in either nucleotide state affects BBSome targeting to the basal bodies. However, the absence of RAB18-GTP prevents BBSome passage through the TZ into cilia (Fig. 3*C* and *D*).

Our previous study demonstrated that the BBSome detaches from retrograde IFT trains in the proximal ciliary region, just above the TZ, and exits cilia via the RABL2-ARL3-mediated outward BBSome TZ diffusion pathway (18, 22, 39). Consequently, the RAB18Q67L mutant most likely causes BBSome buildup in the proximal ciliary region by disrupting its coupling with anterograde IFT trains. TIRF microscopy showed that replacing RAB18 with RAB18-HA did not affect BBSome IFT (Fig. 3*E*, *SI Appendix,* Fig. S6*A* and Movies S10-S11) (21). RAB18S22N-HA failed to restore BBSome IFT (Fig. 3*E*, *SI Appendix,* Fig. S6*A* and Movies S12), as the ciliary BBSome abundance in this strain was likely far below the TIRF detection limit (Fig. 3*A*). Similarly, RAB18Q67L-HA did not restore BBSome IFT (Fig. 3*E*, *SI Appendix,* Fig. S6*A* and Movie S13). This is because the majority of the BBSome remained confined in the proximal ciliary region (Fig. 3*C*) after entering cilia with the aid of the RABQ67L mutant (Fig. 3*A*). Therefore, the GTP-locked RAB18 mutant impedes BBSome integration into anterograde IFT trains in the proximal ciliary region.

### The BBSome acts as a RAB18 effector at the basal bodies

Given that RAB18-GTP promotes BBSome passage through the TZ for ciliary entry, it is likely that RAB18 initially interacts with the BBSome at the basal bodies to facilitate this process. This hypothesis is supported by co-sedimentation of RAB18-HA-YFP (even a small partial) and RAB18Q67L-HA-YFP with the BBSome in sucrose density gradients from the cell body of *RAB18^miR^; RAB18:HA:YFP-TG* and *RAB18^miR^; RAB18Q67L:HA:YFP-TG* cells (Fig. 4*A*). Conversely, RAB18S22N-HA-YFP remained separated from the BBSome in the cell body of *RAB18^miR^; RAB18S22N:HA:YFP-TG* cells (Fig. 4*A*). Our previous study showed that the IFT-A, IFT-B1, IFT-B2, and BBSome subcomplex components of IFT trains remain separated from one another in the HMDEKN buffer (20). In this same buffer, the addition of excessive GTPγS enabled RAB18-HA-YFP to immunoprecipitate the BBSome, but not IFT-A, IFT-B1, and IFT-B2 from the cell body of *RAB18^miR^; RAB18:HA:YFP-TG* cells (Fig. 4*B*). In contrast, excessive GDP addition did not recover any of these complexes, revealing that RAB18 interacts with the BBSome only in its GTP-bound state in the cell body (Fig. 4*B*). Supporting this, RAB18Q67L-HA-YFP, but not RAB18S22N-H-YFP, exclusively recovered the BBSome (Fig. 4*B*). In *E. coli*-based *in vitro* protein interaction assays, RAB18Q67L efficiently captured BBS8, identifying BBS8 as the subunit through which RAB18-GTP directly interacts with the BBSome (Fig. 4*C*). Given that the BBSome and RAB18 colocalize at the basal bodies (Fig. 3*C* and *SI Appendix,* Fig. S4) and the BBSome fails to pass the TZ in the absence of RAB18 or in the presence of RAB18-GDP (Fig. 3*D* and *SI Appendix,* Fig. S4), the interaction between RAB18-GTP and the BBSome as a RAB18 effector likely initiates at the basal bodies.

**Figure 4.**
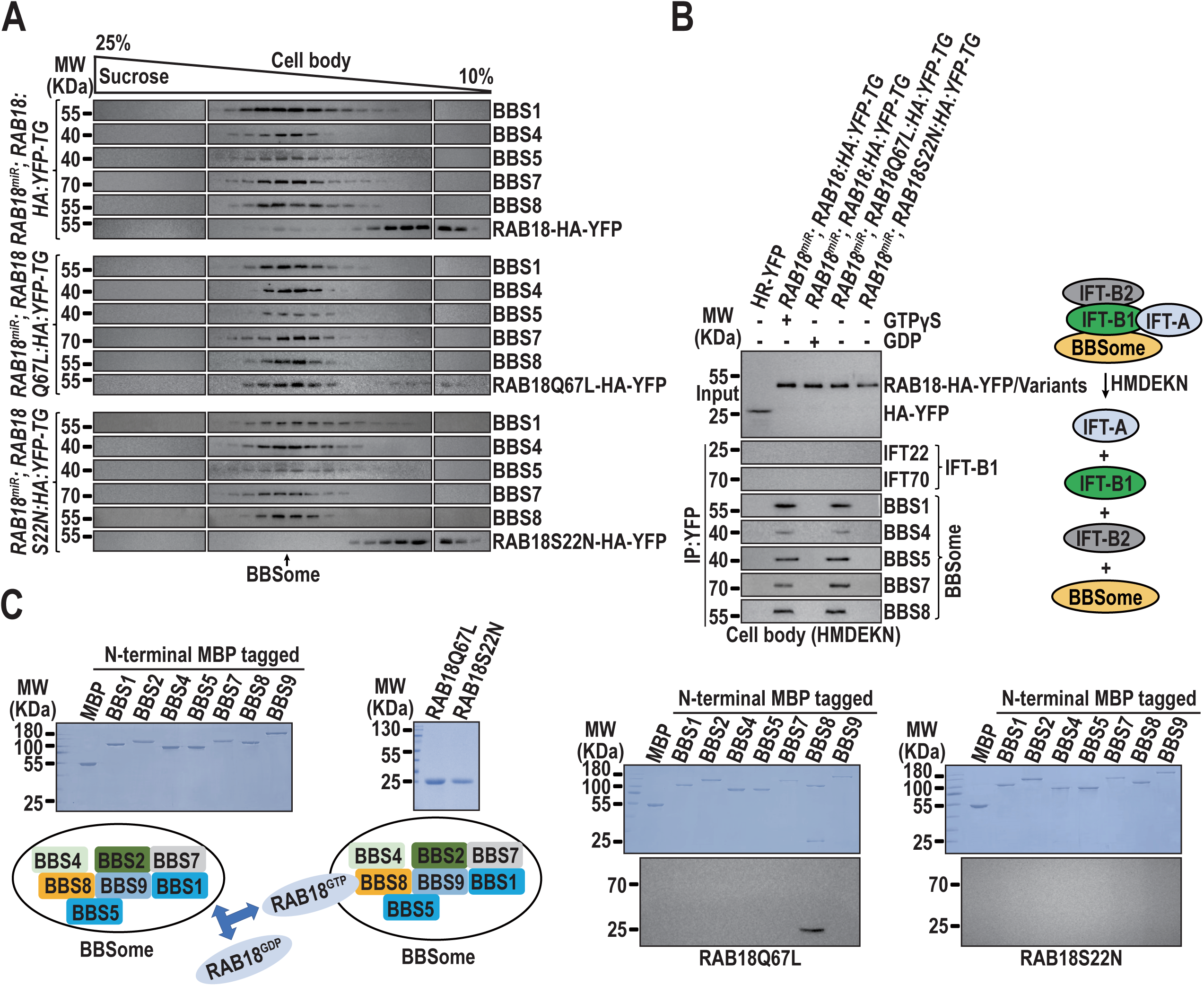
The BBSome acts as a RAB18 effector in a basal body region right below the TZ. **(A)**. Immunoblots of sucrose density gradient fractions of the cell body extracts of the indicated cells probed with α-BBS1, α-BBS4, α-BBS5, α-BBS7, α-BBS8, and α-RAB18. **(B)**. Immunoblots of α-YFP-captured proteins from various cell body extracts: HR-YFP (HA-YFP-expressing CC-125 cells), *RAB18^miR^; RAB18:HA:YFP-TG* (in the presence of excess GTPγS or GDP), *RAB18^miR^; RAB18Q67L:HA:YFP-TG*, and *RAB18^miR^; RAB18S22N:HA:YFP-TG*. Captured complexes were probed for IFT-B1 subunits (IFT22 and IFT70) and BBSome subunits (BBS1, BBS4, BBS5, BBS7, and BBS8). The input was normalized using α-YFP immunoblotting. A schematic on the right illustrates how the BBSome, IFT-B1, IFT-B2, and IFT-A independently exist in HMDEKN buffer. **(C)**. Bacterially expressed MBP, N-terminal MBP tagged BBSome components (BBS1, BBS2, BBS4, BBS5, BBS7, BBS8, and BBS9) (right) were mixed with RAB18Q67L or RAB18S22N (second to the left). Complexes captured on amylose beads were resolved by SDS-PAGE and analyzed by Coomassie staining and immunoblotting with α-RAB18 (second to the right and right, respectively). A schematic at the lower left illustrates the direct interaction between RAB18Q67L and BBS8 of the BBSome. MW, molecular weight.

### The BBSome diffuses through the TZ to enter cilia in an IFT-independent manner

As demonstrated earlier, RAB18-GTP promotes the movement of the BBSome across the TZ for ciliary entry (Fig. 3). To further investigate the interplay between RAB18 and the BBSome during this process, we examined whether the BBSome mediates the ciliary entry of RAB18. To address this question, we generated three strains: *RAB18^miR^; bbs8; RAB18:HA:YFP-TG*, *RAB18^miR^; bbs8; RAB18Q67L:HA:YFP-TG*, and *RAB18^miR^; bbs8; RAB18S22N:HA:YFP-TG*. In these strains, the absence of the BBSome in cilia was ensured by disrupting BBSome assembly through a BBS8 knockout (Fig. 5*A*) (20, 21). When expressed at levels comparable to endogenous RAB18 in CC-5325 cells, all three tagged RAB18 proteins entered cilia through diffusion and maintained at normal levels in cilia, as expected (Fig. 5*A* and *B* and Movies S14- S16, also see Fig. 1*E* and 2*C* and Movies S1-S3). Their ciliary entry, distribution, and fraction patterns was consistent with those observed in *RAB18^miR^* cilia (Fig. 5*B* and *C*; also see Fig. 1*F*, 2*D* and *E*), indicating that the BBSome does not mediate RAB18 ciliary entry.

**Figure 5.**
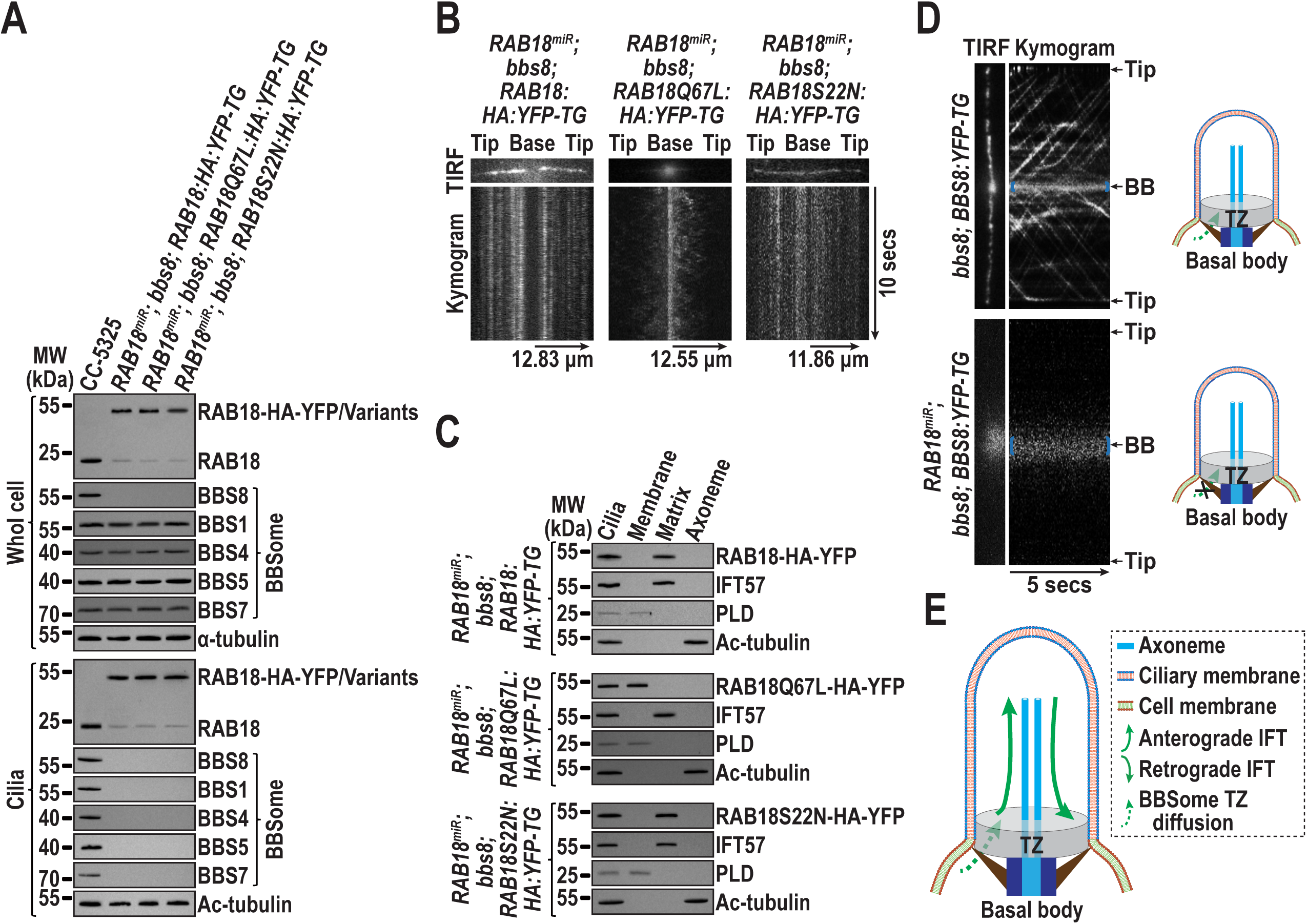
The BBSome diffuses through the TZ for ciliary entry in an IFT-independent manner. **(A)**. Immunoblots of whole cell samples and cilia from cells indicated at the top. Probes include α-RAB18, α- BBS8, α-BBS1, α-BBS4, α-BBS5, and α-BBS7. Alpha-tubulin and Ac-tubulin were used to normalize loading for whole cell samples and cilia, respectively. MW: molecular weight. **(B)**. Representative TIRF images and corresponding kymograms of cells indicated at the top (Movies S14-S16, 15 fps). Time and transport lengths are shown on the right and on the bottom, respectively. The ciliary base (base) and tip (tip) are labeled. **(C)**. Immunoblots of ciliary fractions from cells indicated on the left. Probes include α- YFP, α-IFT57 (ciliary matrix marker), α-PLD (ciliary membrane marker) and Ac-tubulin (axoneme marker). **(D)**. Representative TIRF images and corresponding kymograms of cells indicated on the left (Movies S17-S18, 15 fps). Blue braces highlight the basal bodies. The time is indicated on the bottom. The ciliary base (base) and tip (tip) and the basal body (BB) are shown. A schematic representation illustrates how BBS8-YFP relies on RAB18 for diffusing through the TZ for ciliary entry. **(E)**. Schematic representation of the BBSome’s IFT-dependent diffusion through the TZ for ciliary entry. Once inside, the BBSome integrates into anterograde IFT trains in a proximal ciliary region above the TZ and cycles within cilia via IFT.

RAB18-GTP undergoes lateral transport between the plasma and ciliary membranes, allowing it to diffuse through the TZ for ciliary entry. RAB18-GTP binds to and recruits the BBSome to pass the TZ for ciliary entry, suggesting that the BBSome enters cilia through the same lateral transport mechanism as RAB18-GTP. To test this hypothesis, we knocked down RAB18 to approximately 93% below the endogenous level in *bbs8; BBS8:YFP-TG* cells, resulting in the strain *RAB18^miR^; bbs8; BBS8:YFP-TG* (*SI Appendix,* Fig. S5*C*) (21). TIRF microscopy, as expected, visualized the BBSome (represented by BBS8- YFP) to undergo normal IFT in *bbs8; BBS8:YFP-TG* cilia (Fig. 5*D*, *SI Appendix,* Fig. S6*B* and Movie S17, also see Fig. 3*E* and Movie S10). However, RAB18 knockdown significantly impeded BBS8-YFP entry into cilia to an extend below TIRF detection limit, as evidenced by BBS8-YFP localization at the basal bodies while losing its IFT pattern (Fig. 5*D*, *SI Appendix,* Fig. S6*B* and Movie S18). Directly capturing BBS8-YFP diffusion through the TZ for ciliary entry proven challenging, likely due to the thinness of the TZ, which limits imaging resolution (Fig. 3*E* and 5*D* and Movies S10-S13, S17-S18). However, our data reveal that: 1) RAB18 does not impact IFT (Fig. 1*A-D*); 2) RAB18-GTP diffuses through the TZ for ciliary entry via the membrane system (Fig. 2); 3) RAB18-GTP binds to and recruits the BBSome to move across the TZ and enters cilia (Fig. 3 and 4); and 4) the BBSome remains dissociated from IFT in both the cytoplasma and cilia in the presence of RAB18-GTP (Fig. 4*B* and 6*B*). These results together suggest that the BBSome diffuses through the TZ for ciliary entry in an IFT-independent manner (Fig. 5*E*).

**Figure 6.**
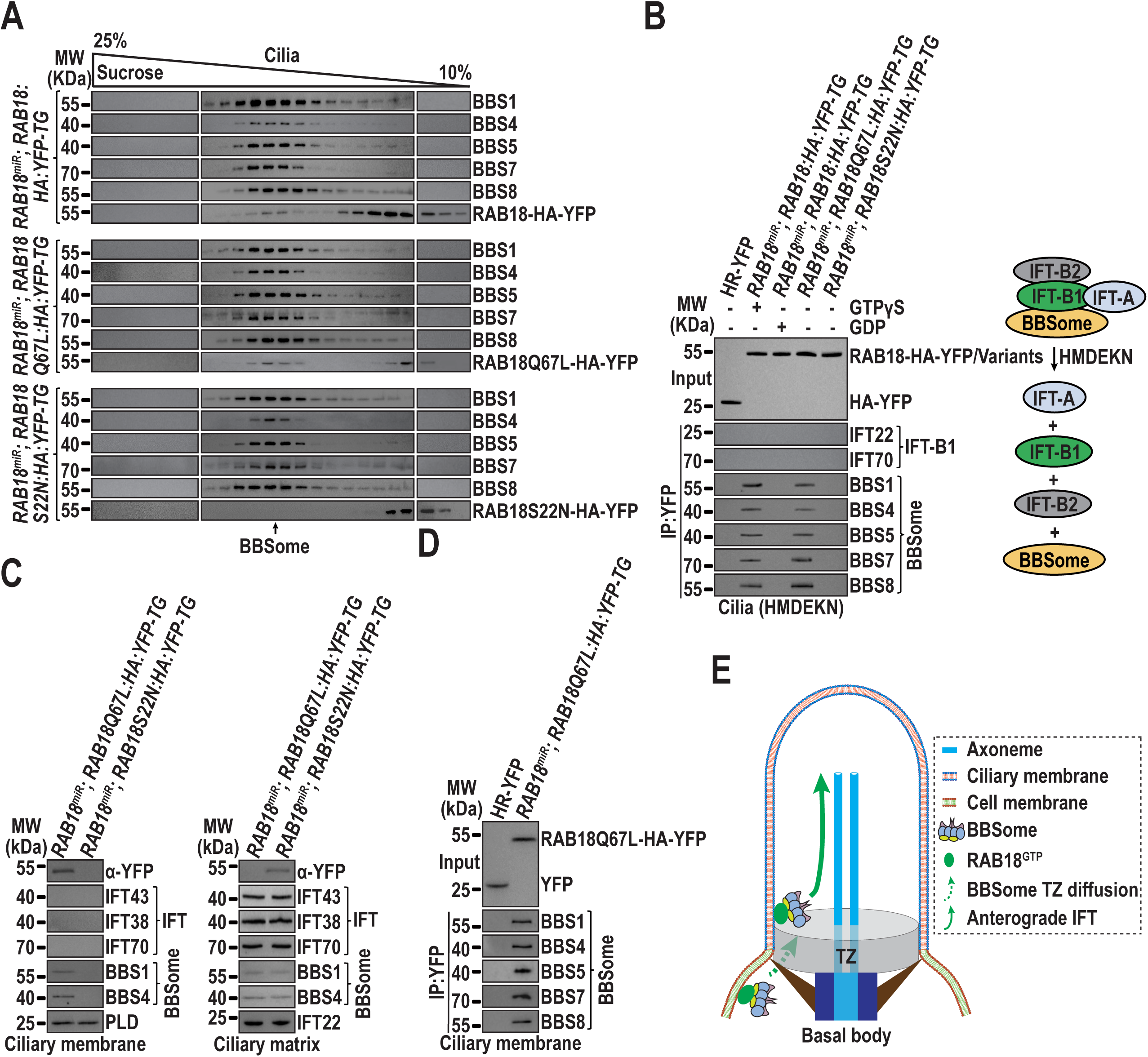
BBSome integrates into IFT trains in a proximal ciliary region. **(A)**. Immunoblots of sucrose density gradients from cilia of the indicated cells probed with α-BBS1, α-BBS4, α-BBS5, α- BBS7, α-BBS8, and α-RAB18. **(B)**. Immunoblots of α-YFP-captured proteins from HR-YFP (HA-YFP- expressing CC-125 cells), *RAB18^miR^; RAB18:HA:YFP-TG* (in the presence of excessive GTPγS or GDP), *RAB18^miR^; RAB18Q67L:HA:YFP-TG*, and *RAB18^miR^; RAB18S22N:HA:YFP-TG* cilia probed for the IFT-B1 subunits IFT22 and IFT70, as well as the BBSome subunits BBS1, BBS4, BBS5, BBS7, and BBS8. HMDEKN is the buffer used for solving cilia. Input was quantified with α-YFP by immunoblotting. A schematic showing the separation of BBSome, IFT-B1, IFT-B2, and IFT-A in HMDEKN buffer is shown on the right. **(C)**. Immunoblots of ciliary membrane (left) and matrix (right) samples from the indicated cells probed for RAB18-HA-YFP variants with α-YFP, the IFT proteins IFT43 (IFT-A subunit), IFT70 (IFT-B1 subunit), and IFT38 (IFT-B2 subunit) and the BBSome subunits BBS1 and BBS4. PLD and IFT22 was used for adjusting the loading of ciliary membrane and matrix, respectively. **(D)**. Immunoblots of α-YFP-captured proteins from *RAB18^miR^; RAB18Q67L:HA:YFP-TG* ciliary membrane probed for the BBSome subunits BBS1, BBS4, BBS5, BBS7, and BBS8. HA-YFP was used as an input and quantified with α-YFP by immunoblotting. **(E)**. Schematic representation of how RAB18-GTP, by anchoring to the ciliary membrane, binds and recruits the BBSome to diffuse through the TZ for ciliary entry in an IFT- independent pattern. The BBSome then integrates into IFT trains in the proximal ciliary region, just above the TZ, and undergoes anterograde IFT. For panels **A-D**, MW stands for molecular weight.

### The BBSome integrates into anterograde IFT trains in the proximal ciliary region

Within cilia, RAB18-HA-YFP partially co-sedimented with the BBSome in sucrose density gradients of *RAB18^miR^; RAB18:HA:YFP-TG* cilia, so did with RAB18Q67L-HA-YFP in *RAB18^miR^; RAB18Q67L:HA:YFP-TG* cilia (Fig. 6*A*). RAB18S22N-HA-YFP inhibited BBSome entry into cilia, however, a trace amount of the BBSome still can enter cilia owing to the RAB18 KD background in *RAB18^miR^; RAB18S22N:HA:YFP-TG* cells (Fig. 3*A*). This allows us to determine whether RAB18S22N-HA-YFP co-sediments with the BBSome in *RAB18^miR^; RAB18S22N:HA:YFP-TG* cilia but the result was negative (Fig. 6*A*). Therefore, the BBSome may remain bound to RAB18-GTP immediately after diffusing into cilia and separates from RAB18 once RAB18-GTP is hydrolyzed into RAB18-GDP. Further supporting this notion, the addition of excessive GTPγS enabled RAB18-HA-YFP to immunoprecipitate the BBSome, but not IFT-A, IFT-B1, and IFT-B2 in *RAB18^miR^; RAB18:HA:YFP-TG* cilia (Fig. 6*B*). Conversely, none of these components was recovered in the presence of excessive GDP (Fig. 6*B*). Consistently, RAB18Q67L-HA-YFP, but not its S22N version, exclusively co-precipitated the BBSome (Fig. 6*B*). We next isolated membrane and matrix contents of *RAB18^miR^; RAB18Q67L:HA:YFP-TG* and *RAB18^miR^; RAB18S22N:HA:YFP-TG* cilia. Both RAB18Q67L-HA-YFP and the BBSome were detected in the ciliary membrane fraction, whereas IFT proteins were absent from this fraction (Fig. 6*C*). On the other hand, the ciliary matrix fraction contained RAB18S22N-HA-YFP, the BBSome, and IFT proteins (Fig. 6*C*). Immunoprecipitation assays performed on the membrane fraction of *RAB18^miR^; RAB18Q67L:HA:YFP-TG* cilia showed that RAB18Q67L-HA-YFP recovered the BBSome, indicating that RAB18-GTP remains bound to the BBSome in the membrane fraction before RAB18-GTP hydrolysis releases the BBSome into the matrix for integration into anterograde IFT trains (Fig. 6*D*). These findings collectively identify an intermediate RAB18-GTP- BBSome complex in the ciliary membrane. TIRF microscopy successfully visualized the BBSome initiating anterograde IFT at the ciliary base, just above the TZ, only in the presence of RAB18-HA-YFP (Fig. 3*D*, 5*D* and Movies S10-S13, S17-S18). Given that the BBSome engages in IFT only after entering cilia, its integration into anterograde IFT trains must occur in the proximal ciliary region immediately above the TZ (Fig. 6*E*).

## Discussion

As a Rab small GTPase, *Chlamydomonas* RAB18 relies on GTP for membrane association, enabling its diffusion into cilia by anchoring to the membrane. The BBSome acts as an effector of RAB18 at the basal bodies, diffusing through the diffusion barrier TZ for ciliary entry via RAB18 (Fig. 7). Our findings demonstrate that RAB18 maintains BBSome ciliary homeostasis by promoting its diffusion across the TZ for ciliary entry via lateral transport between the plasma and ciliary membranes, addressing a critical gap in understanding how RAB18 mediates the export of BBSome cargoes (e.g. PLD) out of cilia in *C. reinhardtii*.

**Figure 7.**
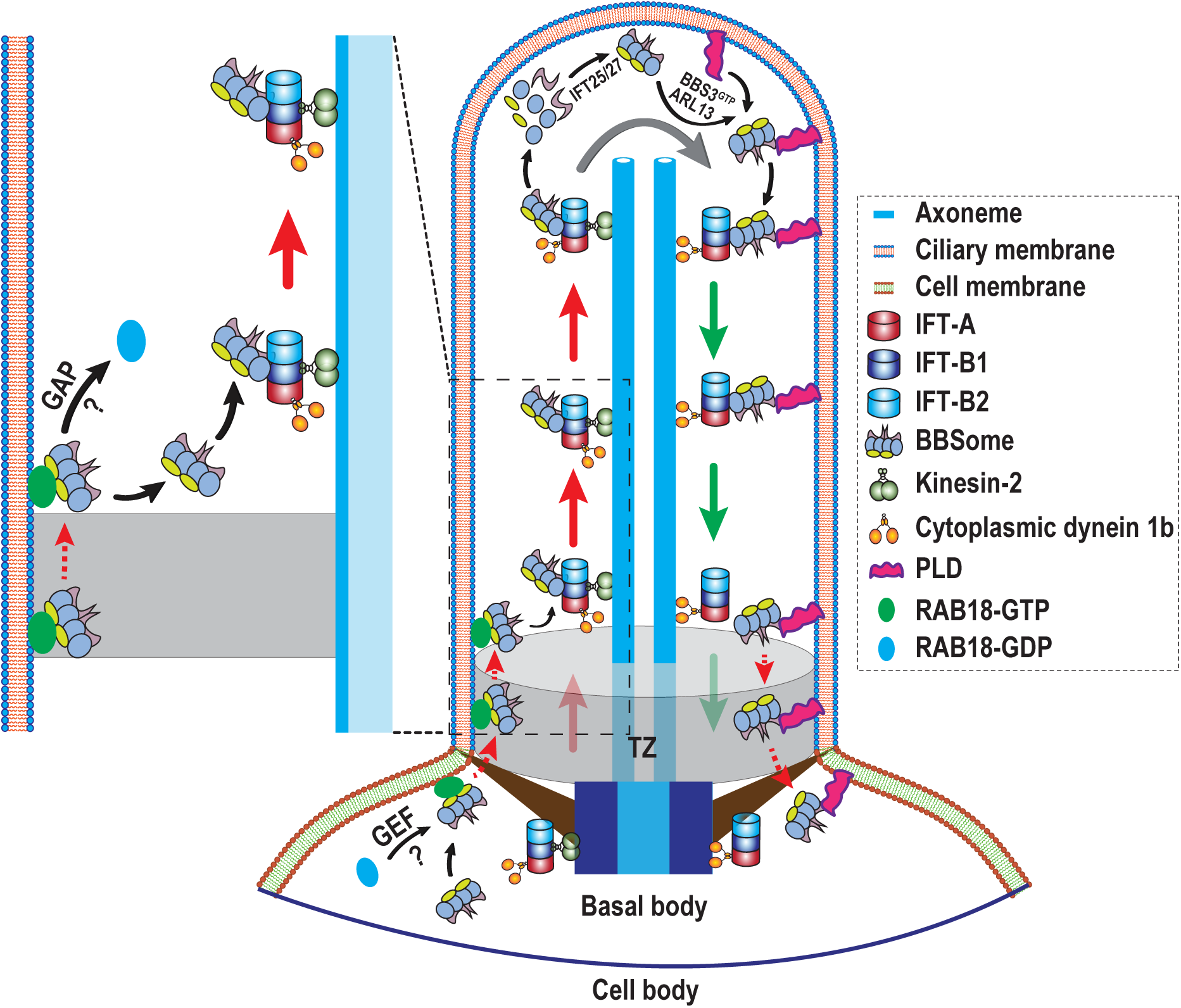
Hypothetical model for how *Chlamydomonas* RAB18 promotes BBSome diffusion through the TZ for ciliary entry. In th basal body region just below the TZ, RAB18 in its GDP-bound form is activated to the GTP-bound form by an unknown GEF (?). The GTP-bound RAB18 is anchored to the cell membrane, where it binds and recruits its effector, the BBSome, autonomously from IFT association, enabling the BBSoem to diffuse through the TZ for ciliary entry, likely through lateral transportation between cell membrane and ciliary membrane. Upon the hydrolysis of the guanine nucleotide by a yet unidentified GAP (?), the BBSome is released from the ciliary membrane and integrates into anterograde IFT trains in the proximal ciliary region just above the TZ. Once at the ciliary tip, the BBSome dissociates from anterograde IFT trains and resides in the ciliary tip matrix, where it remodels with the assistance of IFT-B1-shed IFT25/27 to promote its reassembly (20, 24). The post-remodeled BBSome then couples with its cargo, PLD, via the BBS3-ARL13 module (18, 22). This enables the PLD-laden BBSome to U turn at the ciliary tip and be transported to the ciliary base via retrograde IFT (22). Upon reaching the proximal ciliary region above the TZ, the PLD-laden BBSome dissociates from retrograde IFT trains and diffuses through the TZ for ciliary retrieval via the RABL2-ARL3 module (21, 23).

### Mechanisms of RAB18-mediated BBSome homeostasis in cilia

The BBSome undergoes a seven-step cycling process through cilia across various species: targeting to the basal bodies, crossing the TZ for ciliary entry, anterograde trafficking (from the ciliary base to the tip), ciliary tip remodeling, turnaround at the ciliary tip, retrograde trafficking (from the ciliary tip to the base), and crossing the TZ for ciliary retrieval. This intricate process ensures BBSome ciliary homeostasis and regulates the presence and quantity of signaling proteins within the ciliary membrane, supporting proper cellular signaling. The TZ acts as a diffusion barrier to compartmentalize cilia, facilitating dynamic changes in ciliary composition via regulated trafficking (25, 40, 41). This enables the controlled translocation of large protein complexes, such as the BBSome, across this region (17, 19, 22, 23). In this study, we identified the BBSome as a RAB18 effector at the basal bodies (Fig. 4). RAB18 in its GTP-bound state is anchored to the cell membrane at the basal bodies (Figs. 2 and 3). RAB18-GTP autonomously binds and recruits the BBSome, independent of IFT, for diffusion through the TZ and ciliary entry via lateral transport between the plasma and ciliary membranes (Figs. 3 and 5). While the conservation of this mechanism in mammalian and human RAB18 remains uncertain, our findings suggest that in *C. reinhardtii*, RAB18 ensures BBSome ciliary homeostasis by controlling its availability for TZ-mediated ciliary entry.

### RAB18 likely undergoes nucleotide exchange at the basal bodies

During BBSome ciliary cycling, the BBSome serves as an effector with different GTPases in a compartment-dependent manner. For instance, the BBSome functions as an ARL6 (BBS3) effector during trafficking to the basal bodies (9, 16), as an ARL13 effector at the ciliary tip (22), and as an ARL3 effector in the proximal ciliary region above the TZ (21). Here, we identify the BBSome as a RAB18 effector at the basal bodies, establishing RAB18’s novel role in exporting ciliary membrane proteins (e.g. PLD) by promoting BBSome movement across the TZ for ciliary entry (Fig. 7). Interestingly, RAB18 appears to use the BBSome as an effector only at the basal bodies, not in the cytoplasma or cilia (Fig. 3 and 4). This localized interaction likely prevents competition with IFT22/BBS3, which recruits the BBSome to the basal bodies in its GTP-bound state (16). It is plausible that RAB18 predominantly remains GDP-bound in the cytoplasma, avoiding premature interaction with the BBSome (Fig. 4*A*). Upon reaching the basal bodies, RAB18 undergoes nucleotide exchange to the GTP-bound state, enabling recruitment of the BBSome for TZ diffusion and ciliary entry (Fig. 7). Whether RAB18-GDP requires a GEF for this conversion is unknown, but likely, given the need for precise regulation at the basal bodies (Fig. 7). A similar strategy applies to ARL3 and ARL13, which, target the basal bodies in their GDP-bound state before diffusing into cilia, likely to avoid competing with IFT22/BBS3 for binding to the BBSome in the cytoplasma and basal bodies (21, 22).

### Ciliary base RAB18-GTP is hydrolyzed to release the BBSome for integrating into anterograde IFT trains

Although the BBSome was assumed to integrate into IFT trains at the basal bodies before entering cilia via IFT, our findings and others do not support this hypothesis (42). Once RAB18-GTP diffuses through the TZ for ciliary entry, hydrolysis to RAB18-GDP facilitates BBSome release from the ciliary membrane into the ciliary matrix. This transition is crucial for BBSome integration into anterograde IFT trains in the proximal ciliary region. Although real-time imaging of BBSome diffusion across the TZ was not captured, several biochemical observations strongly support this process: 1) RAB18-GTP binds the membrane, enabling its diffusion through the TZ into cilia; 2) RAB18-GTP recruits the BBSome to cross the TZ for ciliary entry without affecting IFT; and 3) an intermediate RAB18-GTP-BBSome complex exists in the ciliary membrane (Fig. 7). These findings suggest that BBSome integration into anterograde IFT trains occurs only after RAB18-GTP hydrolysis, releasing the BBSome from the ciliary membrane to the ciliary matrix (Fig. 7). Whether RAB18-GTP requires a GAP for this hydrolysis remains unknown, but such a mechanism is plausible in *C. reinhardtii* (Fig. 7).

### Implications for BBS phenotype and RAB18 mutations

In *C. reinhardtii*, the ciliary membrane-anchored PLD associates with the BBSome and diffuses through the TZ for ciliary retrieval via the RABL2-ARL3 module-mediated outward BBSome TZ passage pathway (21, 23). RAB18 promotes inward BBSome TZ passage, regulating its availability for PLD coupling and subsequent ciliary removal (17, 19, 22, 23). If this mechanism applies to human RAB18, its dysfunction caused by certain RAB18 mutations could impair ciliary signaling pathway, such as hedgehog singling, implicated in BBS (43–45). Understanding the molecular interactions between RAB18 and the BBSome offers valuable insights into the pathogenesis of BBS and provides a basis for future research into targeted therapeutic interventions. Further studies are necessary to elucidate the specific effect of RAB18 mutations on BBSome ciliary dynamics and their role in human BBS.

## Materials and Methods

### Antibodies, Chlamydomonas strains, and culture conditions

Rabbit-raised polyclonal antibodies against IFT22 (16), IFT38 (20), IFT43 (46), IFT46 (47), IFT57 (47), IFT70 (47), IFT139 (47), BBS1 (16), BBS4 (18), BBS5 (16), BBS7 (18), BBS8 (20), PLD (18), and CEP290 (21) have been reported previously. Commercially available antibodies used in this study include α-GFP (mAbs 7.1 and 13.1, Roche), α-α-tubulin (mAb B512, Sigma-Aldrich), α-Ac-tubulin (mAb 6-11B- 1, Sigma-Aldrich), as well as goat anti-mouse and anti-rabbit IgGs conjugated to horseradish peroxidase (The Jackson Labs), Alexa-Fluor594, or Alexa-Fluor488 (Molecular Probes). The custom-produced and rabbit-raised α-RAB18 polyclonal antibody was produced by Beijing Protein Innovation LLC (Beijing, China) using bacterially expressed full-length RAB18 as antigen (*SI Appendix*, Fig. S3*A*). A complete list of antibodies utilized in this study is provided in *SI Appendix,* Table S1. Strains CC-5325 and *rab18-876* (LMJ.RY0402.132876) were bought from *Chlamydomonas* Resource Center (https://www.chlamy.org). Previously reported strains include *bbs8* (48); *bbs8; BBS8:YFP-TG* (21), *RAB18; IFT43:HA:YFP-TG* (21), *RAB18; IFT22:HA:YFP-TG* (21), *RAB18; IFT38:YFP-TG* (21), and HA-YFP (49). A complete list of *Chlamydomonas* strains employed in this study is provided in *SI Appendix,* Table S2. *Chlamydomonas* strains were cultured at room temperature in Tris acetic acid phosphate (TAP) medium or minimal 1 (M1) medium under continuous light and constant aeration. Culture conditions varied based on specific strains, with media supplemented as needed with 20 µg/ml paromomycin (Sigma-Aldrich), 15 µg/ml bleomycin (Invitrogen), 10 µg/ml hydromycin (Invitrogen), a combination of two antibiotics (10 µg/ml paromomycin and 5 µg/ml bleomycin), or three antibiotics (10 µg/ml paromomycin, 5 µg/ml bleomycin, and 10 µg/ml hydromycin).

Various experimental protocols were employed in this study and are briefly described in the text for clarity. Detailed descriptions of all protocols are provided in *SI Appendix*.

### Statistics

Statistical analyses were performed using GraphPad Prism 8.0 (GraphPad Software). Data are presented as mean ± S.D. from three independent experiments. The sample size is denoted by “n.” A one-sample unpaired Student’s *t*-test was used to compare velocities and frequencies of YFP-tagged proteins. Two-tailed *t*-tests were employed to analyze protein expression levels and ciliary lengths. Results marked as “n.s.” indicate non-significant differences.

### Data, Materials, and Software Availability

All study data are included in the article and/or supporting information. Materials used in this study are available upon request.

## Acknowledgements

Research reported in this publication was supported by the National Natural Science Foundation of China (32470741 to Z.-C.F.). We thank the stuff from the Core Facilities, the Fourth Affiliated Hospital of School of Medicine, and International School of Medicine, International Institutes of Medicine, Zhejiang University, for their technical support. The founder has no role in study design, data collection and analysis, decision to publish, or preparation of the manuscript.

## Author contributions

Z.-C.F. designed research; W.-Y.S., J.Z., Y.W. and L.S.Z. performed research; W.-Y.S. and Z.-C.F. analyzed data; and Z.-C.F. wrote the paper.

## Competing financial interests

The authors declare no conflict of interest.

## Materials and methods

### Plasmids

The miRNA targeting 3’-UTRs of the *rab18* gene was designed with the WMD3 software (http://wmd3.weigelworld.org) (1). The output oligonucleotide was combined with the miRNA cre- MIR1157 (accession number MI0006219), resulting in the miRNA precursor sequence of 171-bp for *rab18* (*SI Appendix*, Table S3). The sequence was synthesized by the commercial company Genewiz (China) and ligated to the *Eco*RV and *Eco*RI sites of pHK263 plasmid (1), resulting in pMi- RAB18-Paro.

*Chlamydomonas* expression vectors were constructed on pBKS-gRABL2-HA-YFP-Ble and pBKS- gRABL2-HA-Ble that contained a HA-YFP (pBKS-gRABL2-HA-YFP-Ble) and HA (pBKS-gRABL2- HA-Ble) coding sequence followed immediately downstream by a sequence encoding the Rubisco 3’-UTR and the bleomycin (*Ble*, zeocine resistant gene) cassette (2). Genomic DNA of *Chlamydomonas* cells was extracted and purified using a Wizard® Genomic DNA Purification Kit (Promega, Beijing) following the kit’s protocol. To generate the RAB18-HA-YFP expression vector, a 4,024-bp *rab18* DNA fragment containing 1,000-bp promoter sequence was amplified from genomic DNA by using the primer pair gRAB18-FOR and gRAB18-REV as listed in *SI Appendix*, Table S3. This fragment was inserted into pBluescript II KS(+) vector linearized by *Xba*I and *Eco*RI, resulting in pBKS-gRAB18. Site-directed mutagenesis was performed to introduce the Q67L and S22N mutations into pBKS-gRAB18 by using the primer pairs RAB18Q67L-FOR and RAB18Q67L- REV and RAB18S22N-FOR and RAB18S22N-REV, respectively, as listed in *SI Appendix*, Table S3. These *Xba*I and *Eco*RI double digested RAB18/variant DNA fragments and the HA-YFP-Ble or HA-Ble DNA fragment cut from pBKS-gRABL2-HA-YFP-Ble and pBKS-gRABL2-HA-Ble by *Eco*RI and *Kpn*I were inserted into pBluescript II KS(+) vector linearized by *Xba*I and *Kpn*I by three-way ligation, resulting in pBKS-gRAB18-HA-YFP-Ble, pBKS-gRAB18Q67L-HA-YFP-Ble, and pBKS- gRAB18S22N-HA-YFP-Ble, pBKS-gRAB18-HA-Ble, pBKS-gRAB18Q67L-HA-Ble, and pBKS-gRAB18S22N-HA-Ble, respectively.

The bacterial expression vectors were generated as described previously (3). To make these vectors generated, total RNA of *Chlamydomonas* cells was extracted and purified using Eastep® Super total RNA Extract Kit (Promega, Shanghai). Five micrograms of RNA were reverse transcribed at 42 °C for 1 h using M-MLV Reverse Transcriptase (Promega) and oligo(T)18 primers (Takara). The *rab18* cDNA was amplified by PCR using primer pair cRAB18-FOR and cRAB18- REV as listed in *SI Appendix*, Table S3. The PCR reactions were performed at 95 °C for 5 min followed by 30 cycles of 95°C for 20 sec, 61°C for 20 sec, and 72°C for 4 min. The DNA fragment obtained was inserted into the *Eco*RI and *Xho*I sites of pET-28a (Novagen) for generating pET- 28a-cRAB18. Q67L and S22N mutations were introduced into the *rab18* cDNA by site-directed mutagenesis using primer pairs RAB18Q67L-FOR and RAB18Q67L-REV and RAB18S22N-FOR and RAB18S22N-REV, respectively, as listed in *SI Appendix*, Table S3, resulting in pET-28a- cRAB18Q67L and pET-28a-cRAB18S22N.

All new vectors were verified by direct nucleotide sequencing.

### Transgenic strain generation

The *C. reinhardtii* strain CC-5325 was transformed with pMi-RAB18-Paro for generating the RAB18 miRNA strain *RAB18^miR^*. The *RAB18^miR^* strain was transformed with pBKS-gRAB18-HA-YFP-Ble, pBKS-gRAB18Q67L-HA-YFP-Ble, pBKS-gRAB18S22N-HA-YFP-Ble, pBKS-gRAB18-HA-Ble, pBKS-gRAB18Q67L-HA-Ble, and pBKS-gRAB18S22N-HA-Ble for generating the rescuing strains including *RAB18^miR^; RAB18:HA:YFP-TG*, *RAB18^miR^; RAB18Q67L:HA:YFP-TG*, *RAB18^miR^; RAB18S22N:HA:YFP-TG*, *RAB18^miR^; RAB18:HA-TG*, *RAB18^miR^; RAB18Q67L:HA-TG*, and *RAB18^miR^; RAB18S22N:HA-TG*. DNA transformation was conducted by electroporation according to the previously reported protocol (4) and the transformants were selected on TAP plates with 20 µg/ml paromomycin (Sigma-Aldrich), 15 µg/ml bleomycin (Invitrogen) or both antibiotics with 10 µg/ml paromomycin and 5 µg/ml bleomycin. The screening of RAB18 miRNA cells was initiated by checking the cellular level of the target proteins through immunoblotting of whole cell extracts with RAB18 antibody. The miRNA strains showing a reduced level of the target proteins were selected for further phenotypic analysis. The positive transformants were identified and the target proteins were quantified directly by immunoblotting as described below.

### *Chlamydomonas* strain crossing

*RAB18^miR^; bbs8; RAB18:HA-TG; BBS8-YFP-TG*, *RAB18^miR^; bbs8; RAB18Q67L:HA-TG; BBS8-YFP-TG*, and *RAB18^miR^; bbs8; RAB18S22N:HA-TG; BBS8-YFP-TG* were generated through the crossing of strains *RAB18^miR^; RAB18:HA-TG*, *RAB18^miR^; RAB18Q67L:HA-TG*, and *RAB18^miR^; RAB18S22N:HA-TG* with *bbs8; BBS8:YFP-TG* that has been reported previously (2). In briefly, 20 ml cultures of each strain (*RAB18^miR^; RAB18:HA-TG*, *RAB18^miR^; RAB18Q67L:HA-TG*, *RAB18^miR^; RAB18S22N:HA-TG*, and *bbs8; BBS8:YFP-TG*) were grown in M1 medium until reaching a density of 2×10^6^ cells/ml. Subsequently, cells were washed with M1-N (without nitrogen) medium three times, transferred to 20 ml of M1-N medium, and aerated overnight (∼12 hrs) under constant light at RT. Gametes of two opposite mating types were mixed in flask and incubated under light for 2 hrs. The cells settled at the bottom of the flask were transferred to M1 plate containing 4% agar, air-dried, and then incubated overnight under continuous light. Following this, cells were incubated in the dark for a week before being moved to incubate at -20°C for two days. Subsequently, cells were moved to incubate under continuous light at RT for at least ten days. *RAB18^miR^; bbs8; RAB18:HA-TG; BBS8-YFP-TG*, *RAB18^miR^; bbs8; RAB18Q67L:HA-TG; BBS8-YFP-TG*, and *RAB18^miR^; bbs8; RAB18S22N:HA-TG; BBS8-YFP-TG* were screened by detecting the loss of RAB18 and BBS8 and the presence of RAB18-HA, RAB18Q67L-HA. RAB18S22N-HA and BBS8- YFP proteins through immunoblotting of whole cell samples. The immunoblotting involved the use of antibodies specific to RAB18, BBS8, HA, and YFP, as described below.

### Isolation of cilia and cell bodies

*Chlamydomonas* cilia and cell bodies were isolated by following a previously reported protocol (5). Unless otherwise stated, cells were cultured in 10 liters of TAP medium until reaching a density of 10^8^ cells/ml. The cells were then harvested by centrifugation at 1,000 *×g* for 15 min at room temperature (RT), resuspended in 150 ml TAP medium (pH7.4), and incubated for 2 hrs under strong light with bubbling. Subsequently, 0.5 M acetic acid was added to adjust the pH to 4.5 for deciliation within 30-60 sec. The pH was then restored to 7.4 by adding 0.5 M KOH. *C*ell bodies were pelleted by centrifugation at 600 *×g* 4°C for 5 min. The supernatant was further centrifuged at 12,000 ×*g* 4°C for 10 min, and the resulting pellets were collected as cilia. Cilia were thoroughly washed with HMDEKN buffer (30 mM Hepes [pH 7.4], 5 mM MgSO_4_, 1 mM DTT, 0.5 mM EGTA, 25 mM KCl, 125 mM NaCl) supplemented with protein inhibitors (PI) (1 mM PMSF, 50 μg/ml soy- bean trypsin inhibitor, 1 μg/ml pepstatin A, 2 μg/ml aprotinin, and 1 μg/ml leupeptin) through centrifugation at 12,000 ×*g* 4°C for 10 min until the green color disappeared completely.

### Preparation of ciliary fractions

Ciliary membrane, matrix, and axoneme were isolated using a previously established protocol (6). In brief, cilia suspended in HMDEKN buffer plus PI underwent three cycles of freezing and thawing in liquid nitrogen. After centrifugation at 12,000 ×*g* at 4°C for 15 min, the supernatant was collected as the ciliary matrix. The pellets, consisting of ciliary membrane and axoneme, were resuspended in HMEDKN buffer plus PI supplemented with 1% nonidet P-40 (NP-40) and allowed to sit on ice for 30 min. The supernatant and pellets were separated as ciliary membrane and axonemal fractions, respectively, by centrifugation at 12,000 ×*g* at 4°C for 10 min. For immunoblotting assays, 20 μl of each ciliary fraction samples were loaded into 12% SDS-PAGE gel.

### Sucrose density gradient centrifugation assay

Sucrose density gradient centrifugation of cell body samples and cilia was conducted following our previously reported protocol (7). In brief, linear 12 ml gradients of 10-25% sucrose density were prepared in HMDEKN. Cell bodies or cilia suspended in HMDEKN plus PI supplemented with 1% NP-40 underwent three cycles of freezing and thawing in liquid nitrogen. After centrifugation at 12,000 *×g* at 4 °C for 10 min, the non-soluble debris was removed and 700 μl of supernatants were loaded on top of the gradients. Centrifugation was performed at 38,000 rpm at 4 °C for 14 hrs in a SW41Ti rotor (Beckman Coulter). Subsequently, the gradients were collected in 24 fractions of 0.5 ml aliquots. To detect protein co-sedimentation patterns, 20 μl of each fraction was loaded into 12% SDS-PAGE gel for electrophoresis and analyzed by immunoblotting, as described below. The standards used to calculate S-values were BSA (4.4S), aldolase (7.35S), catalase (11.3S), and thyroglobulin (19.4S).

### Immunoblotting

Immunoblotting was conducted on whole cell, cell body, cilia and ciliary fraction samples, as well as immunoprecipitated protein samples, following our previously reported protocol (8). Samples were mixed with Laemmli SDS sample buffer and boiled for 5 min, followed by centrifugation at 2,500 *×g* for 5 min to remove debris. Unless specified otherwise, 20 µg (whole cell and cell body samples and immunoprecipitated protein samples) or 100 µg (cilia and ciliary fraction samples) of total protein from each sample was loaded onto a 12% SDS-PAGE gel for electrophoresis before transfer to nitrocellulose (NC) membrane. The NC membranes were blocked in 5% non-fat dry milk (Acmec) in TBS (10 mM Tris, pH 7.5, 166 mM NaCl) plus 0.05% Tween-20 and incubated with primary antibodies diluted in the blocking solution for 2 hrs at RT. Following three washes with TBS plus 0.05% Tween-20, the membranes were incubated with HRP-conjugated secondary antibodies for 2 hrs at RT. After five additional washes, chemiluminescence (Millipore) was used to detect the primary antibodies, employing the Mini Chemiluminescent/Fluorescent Imaging and Analysis System (Sagecreation). Primary and secondary antibodies were diluted for immunoblotting according to the ratios shown in *SI Appendix*, Table S1. If necessary, ImageJ software (version 1.42g, National Institutes of Health) was employed to quantify target proteins by measuring the immunoblot intensity, following our previously reported protocol (9). The immunoblot intensity was normalized to the intensity of a loading control protein.

### Immunoprecipitation

Immunoprecipitation was conducted on cell body, cilia, and ciliary fraction samples according to our previously reported protocol (6). Briefly, samples prepared from the cell body, cilia, and ciliary fractions of *Chlamydomonas* cells expressing HA-YFP, HA-YFP-tagged RAB18 and its variants were resuspended in HMDEKN plus PI supplemented with 1% NP-40 and subject to three cycles of freezing and thawing in liquid nitrogen. After centrifugation at 70,000 ×*g* at 4°C for 1 hr, the supernatants were collected and agitated with 5% BSA-pretreated camel anti-YFP antibody-conjugated agarose beads (V-nanoab Biotechnology) for 2 hrs at 4°C. Following centrifugation at 2,500 ×*g* at 4°C for 2 min, the beads were washed three times with 1 ml HMDEKN buffer, twice with 1ml HMDEK buffer containing 50 ml NaCl, and twice with 1ml HMDEKN. The beads were then incubated in 100 μl of 0.1 M Glycine buffer (pH2.5) for 2 min at RT, followed by centrifugation at 2,500 ×*g* at 4°C for 2 min to collect the supernatant. The supernatants were mixed with Laemmli SDS sample buffer and boiled for 5 min before centrifugation at 2,500 *×g* for 5 min. The immunoprecipitants in the supernatants were collected for immunoblotting. If required, 20 mM GTPγS or GDP were included for the whole experimental process.

### Liposome flotation assays

The bacterial expression vectors including pET-28a-cRAB18, pET-28a-cRAB18Q67L, and pET- 28a-cRAB18S22N were transformed into the *Escherichia coli* BL21(DE3) cell and the C-terminal 6*×*His tagged RAB18 or its variants were purified with Ni Sepharose^TM^ 6 Fast Flow beads (GE Healthcare) and, thereafter, the His-tag was removed by thrombin (Solarbio) at the concentration of 3 U/mg as described in our previous report (9). Liposome flotation assays were performed according to our previous report (2). In brief, rehydrated lipid films (Avanti) in the lipid reconstitution buffer (30 mM Tris, 150 mM NaCl, 2 mM MgCl_2_, 2 mM DTT, pH 7.5) were mixed by centrifuging at 1,000 rpm for 1 hr at RT. Liposomes were generated from re-suspended lipids with an extruder (200 nm pore size, 20 strokes manually, Avanti). Thereafter, 70 µg of purified RAB18 (100 μM) was incubated in the presence of 500 μM GTPγS or GDP. MgCl_2_ was added to samples to reach a final concentration of 100 mM and the RAB18 protein was separated from the excess of nucleotides by Zeba Spin desalting columns (Thermal Scientific). Afterward, 75 µl of liposomes and lipid reconstitution buffer were mixed to reach a final volume of 150 µl and incubated for 2 hrs at RT. The reactions were added with 100 µl 75% sucrose solution, 200 µl 25% sucrose, and 50 µl lipid reconstitution buffer, and centrifuged at 240,000 ×g for 1 hr (Beckman TLS-55 rotor, OptiMax MAX-XP benchtop centrifuge). After the centrifugation, 50 µl solutions were collected from the top of the centrifuge tube and subjected to SDS-PAGE electrophoresis and Coomassie staining. The purified RAB18 variants were used for liposome flotation assays without nucleotide preloading. For SDS-PAGE and Coomassie staining, 1 µg of RAB18 or its variant proteins was loaded for evaluating their binding/input ratio shown as a percentile.

### Fixed imaging

Immunofluorescence staining was performed according to our previous report (10). Briefly, after *Chlamydomonas* cells growing in M1 media were washed once with 5 mM EGTA, they were seeded to 0.1% polyethyleneimine-coated coverslips for 8 min. Cells were permeabilized and fixed with ice-cold methanol twice each for 10 min and rehydrated with phosphate buffered saline (PBS). Thereafter, cells were blocked in blocking buffer (5% BSA, 1% cold water fish gelatin, and 10% goat serum in PBS) for 4 hrs and incubated with primary antibodies in blocking buffer for 2 hrs. Cells were then washed ten times in PBS followed by incubating with secondary antibodies in blocking buffer for 1 hr. After washing additional ten times in PBS, the coverslips were mounted with SlowFade Antifade reagent (Molecular Probes) and images were captured with an Olympus SpinSR spinning disk confocal microscopy equipped with a Hamamatsu ORCA-Fusion sCMOS camera (C14440-20UP), a 100×/1.40 NA oil objective lens (Olympus), and 488-nm and 561-nm lasers from Coherent OBIS Laser Module. All images were acquired and processed with CellSens Dimension (version 2.1, Olympus) and shown as grayscale. The primary antibodies against YFP, BBS8, and CEP290 and the secondary antibodies Alexa-Fluor488-conjugated goat anti-mouse and Alexa-Fluor594-conjugated goat anti-rabbit (Molecular Probes) were listed in *SI Appendix*, Table S1 with their suggested dilutions. The staining was performed at RT.

### Total internal reflection fluorescence (TIRF) microscopy and kymogram analysis

Cells expressing YFP-tagged RAB18 and its variants (10 μl) growing in M1 medium were placed onto the larger coverslip (24×60-mm No.1.5) followed by settling for 2 min at RT. A ring of petroleum jelly was then added around the droplet and a smaller coverslip (22×22-mm No.1.5) containing 5 μl of 10 mM HEPES, pH 7.4 and 5 mM EGTA was inverted onto the larger one to generate a sealed observation chamber. TIRF microscopy was applied to visualize the motility of YFP-tagged RAB18 and its variants in cilia. Videos were captured with TIRF (15 Hz) for ∼20 s by using the software Vsim (Nikon) at 15 frames per second (fps). For capturing video, a Nikon Ti2-E inverted fluorescent microscopy equipped with a through-the-objective TIRF system, a 100×/1.49 NA TIRF oil immersion objective len (Nikon), a back illuminated scientific CMOS camera (Kinetix, Photometrics), and 488-nm laser from Coherent OBIS Laser Module was used according to our previous report (9). The videos were processed with ImageJ software (version 1.52f, National Institutes of Health) for generating kymographs according to our protocol reported previously (8).

### Protein-protein interaction assay

Bacterially expressed N-terminal MBP-tagged BBS1, BBS2, BBS4, BBS5, BBS7, BBS8, and BBS9 proteins were purified according to our previous report (2). To determine RAB18 interaction to the BBSome, 100 μg of MBP and the MBP-tagged BBS proteins were individually mixed with 100 µg of bacterially expressed RAB18Q67L or RAB18S22N (see above) to form a combination of 16 reactions. After incubated for 2 hrs at RT, mixtures were purified with Dextrin Sepharose^TM^ High Performance MBP-tagged protein purification resin (GE Healthcare). Ten micrograms of proteins from elutes was loaded on 12% SDS-PAGE gels for electrogenesis. Gels were then visualized with Coomassie blue staining. Immunoblotting assay was also performed to verify the interaction between BBS proteins and RAB18 variants with α-RAB18.

### Measurement of ciliary length

Differential interreference contrast images of *Chlamydomonas* cell were taken by using an Olympus IX73 inverted microscope equipped with a Hamamatsu ORCA-Fusion4.0LT camera (C11440- 42U30) under a 60× objective. Ciliary length was measured according to our previous report (11).

---

**Fig. S1.**
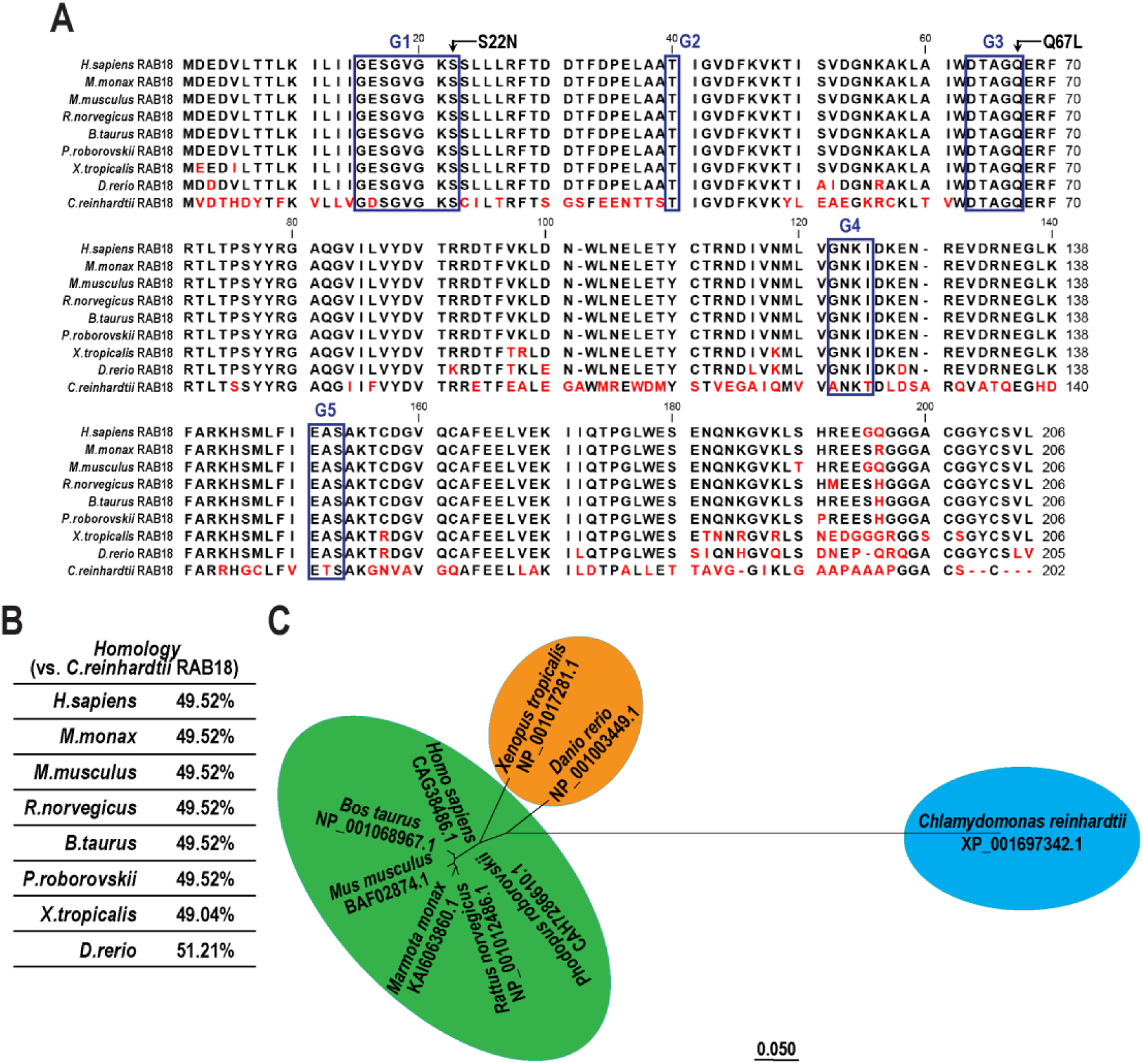
RAB18 is highly conserved across ciliated species. (A). Sequence alignment of deduced amino acid sequences from nine invertebrate and vertebrate RAB18 orthologues. Alignments were generated using CLC main workbench (version 6.8); the most conserved residues are shown in black, the least conserved are in red. RAB18 contains five conserved domains including G1, G2, G3, G4, and G5 as labeled in the boxes. The P-loop threonine (S22) and glutamine (Q67) were marked. Dashes indicate gaps introduced to optimize the alignment. Arrowheads indicate missense mutations created in possible dominant-negative (S22N) and constitutive-active (Q67L) RAB18 mutants. (B). A table shows the identity of *Chlamydomonas* RAB18 in comparison to its homologues in other species as indicated. (C). The phylogenetic tree of RAB18 proteins from invertebrate and vertebrate species as indicated. The neighbor-joining tree was calculated using the MEGA 7 software. Branch length represents evolutionary relatedness. For three panels, GenBank accession numbers are as follows: *Homo sapiens*, CAG38486.1, *Marmota monax*, KAI6063860.1, *Mus musculus*, BAF02874.1, *Rattus norvegicus*, NP_001012486.1, *Bos Taurus*, NP_001068967.1, *Phodopus roborovskii*, CAH7286610.1, *Xenopus tropicalis*, NP_001017281.1, *Danio rerio*, NP_001003449.1 and *Chlamydomonas reinhardtii*, XP_001697342.1.

**Fig. S2.**
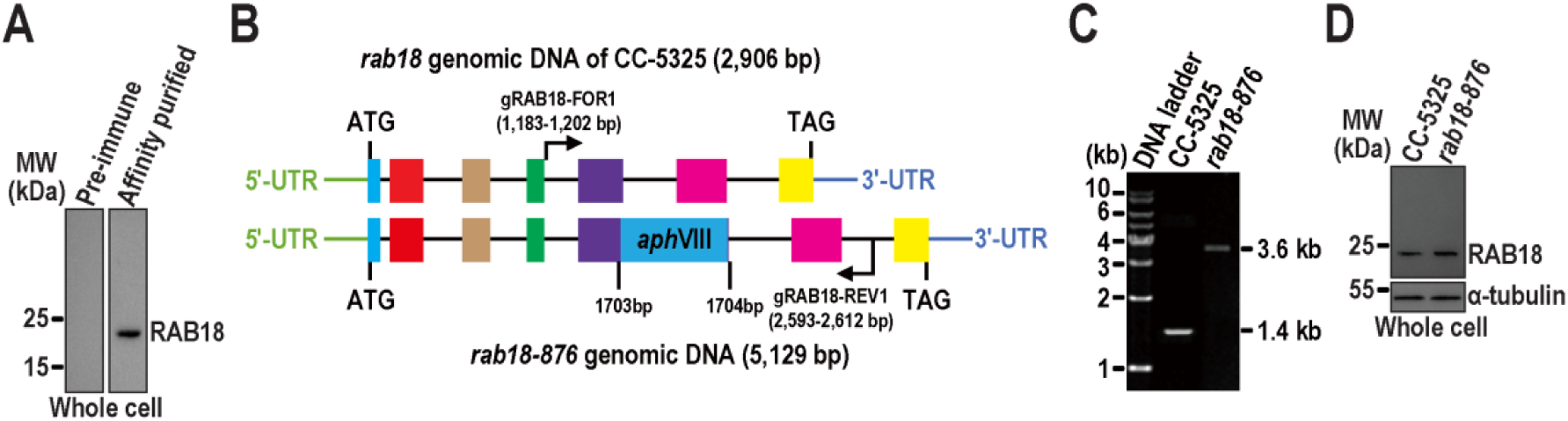
Characterization of the RAB18 CLiP strain named *rab18-876* (LMJ.RY0402.132876). (A). Immunoblots with the affinity-purified RAB18 antiserum but not the pre-immune serum identified RAB18 protein with a size of approximately 20 kDa in whole cell samples of CC-5325. MW stands for molecular weight. (B). A schematic representation shows that a paromomycin-resistant gene (*AphVIII*) inserted at 3’-proximal region of the fifth exon in the *rab18* gene of *rab18-876* cells. The colored boxes represent the exons, and the lines represent the introns of the *rab18* gene, respectively. (C) Agarose gel electrophoresis of the PCR products amplified from genomes of CC- 5325 and *rab18-876* cells. The primer pair gRAB18-FOR1 and gRAB18-REV1 were used to amplify the *rab18* genomic DNA of 1,429-bp from CC-5325 cells. A single DNA fragment of ∼3.6-kb (3,652- bp) was amplified from the *rab18-876* cells by using the same primer pair. (D). Immunoblots of whole cell samples of CC-5325 and *rab18-876* probed with α-RAB18. Alpha-tubulin we used as a loading control. MW stands for molecular weight.

**Fig. S3.**
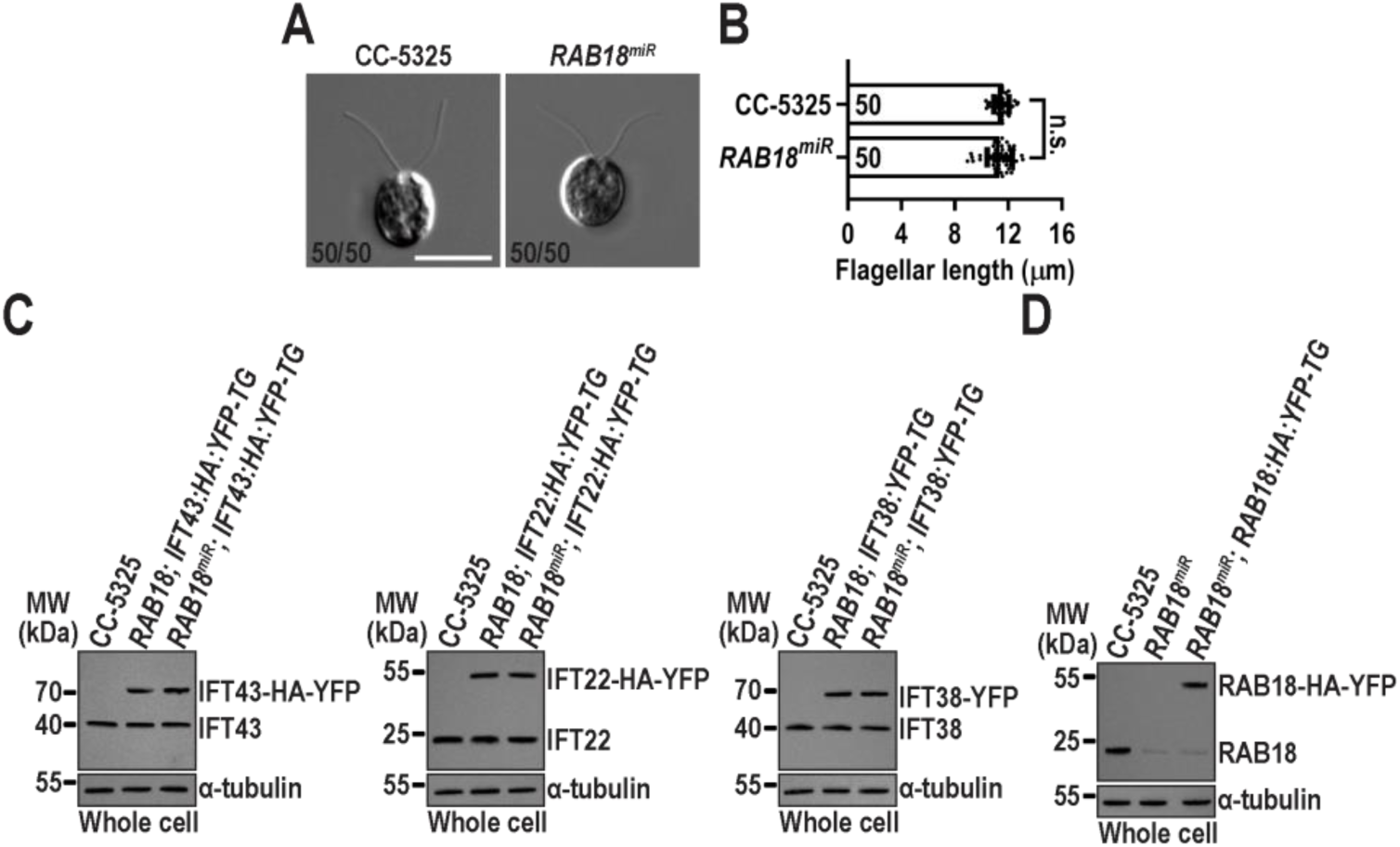
*Chlamydomonas* RAB18 is dispensable for ciliation and IFT. (A). *RAB18^miR^* cells assemble cilia with normal length. Representative differential interreference contrast images of CC-5325 and *RAB18^miR^* cells are shown. The number of cells with cilia out of fifty counted cells is listed for each strain. Scale bar: 10 µm. (B). *RAB18^miR^* cells had full-length cilia (11.49 ± 0.55 µm, n = 50) compared to CC-5325 cells (11.35 ± 0.94 µm, n = 50). Mean lengths are listed; error bar indicates S.D., and “n” indicates the number of cilia counted and listed in each bar. “n.s.” denotes non-significance. A two-tailed *t*-test is used for statistical analysis. (C). Immunoblots of whole cell samples from three cell groups, including CC-5325, *RAB18; IFT43:HA:YFP-TG*, and *RAB18^miR^; IFT43:HA:YFP-TG* (left); CC-5325, *RAB18; IFT22:HA:YFP-TG*, and *RAB18^miR^; IFT22:HA:YFP-TG* (middle); and CC-5325, *RAB18; IFT38:YFP-TG*, and *RAB18^miR^; IFT38:YFP-TG* (right) probed with α-IFT43, α-IFT22, and α-IFT38, respectively. For each group of the cells, the YFP- or HA-YFP-tagged proteins of both CC-125 (*RAB18; IFT43:HA:YFP-TG*, *RAB18; IFT22:HA:YFP-TG*, and *RAB18; IFT38:YFP-TG*) and *RAB18^miR^* background were determined to express at the wild-type CC-5325 protein levels. (D). Immunoblots of whole cell samples from CC-5325, *RAB18^miR^*, and *RAB18^miR^; RAB18:HA:YFP-TG* probed with α-RAB18. For panels C and D, α-tubulin we used as a loading control. MW stands for molecular weight.

**Fig. S4.**
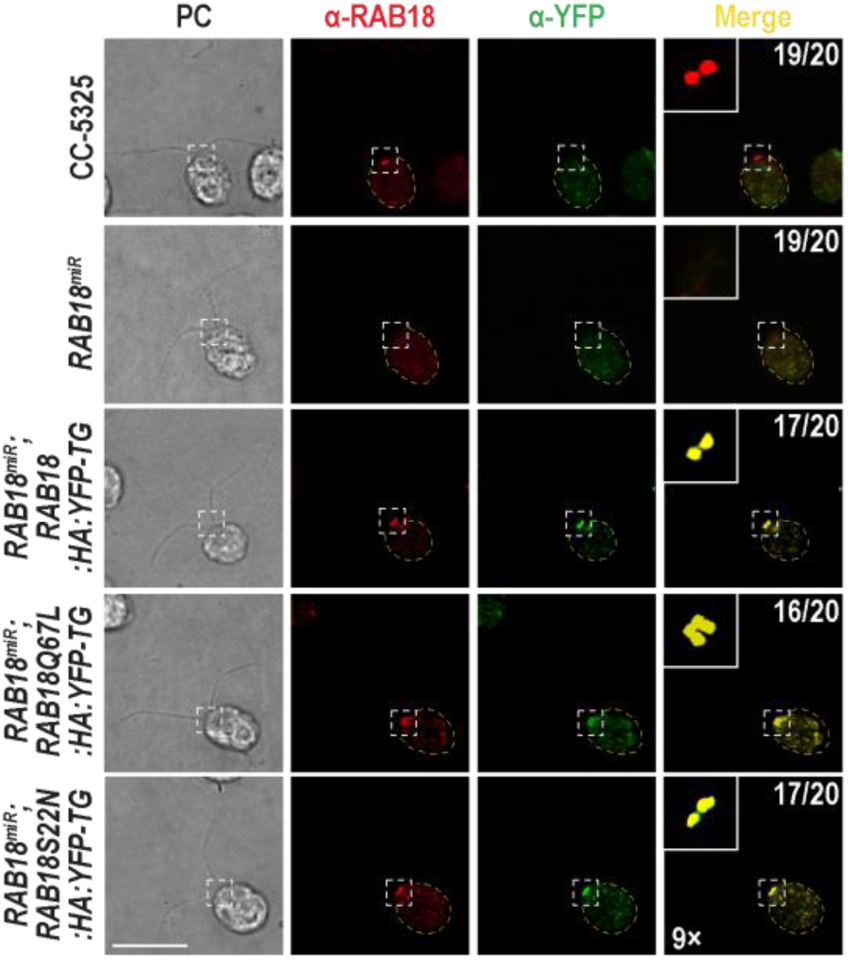
The C-terminal HA and YFP double tagged RAB18 and its variants resemble the endogenous RAB18 in distributing to the basal bodies. Immunofluorescence staining of the indicated cells with α-RAB18 (red) and α-YFP (green). Phase contrast (PC) images of cells are included. Insets highlight the proximal ciliary region above the TZ and the basal bodies. Inset magnifications (9 times) are displayed. Scale bars: 10 µm. Cell numbers out of 20 cells are listed for representing cells that are positive in RAB18 (RAB18-HA-YFP and its variants) distribution to the proximal ciliary region above the TZ and the basal bodies.

**Fig. S5.**
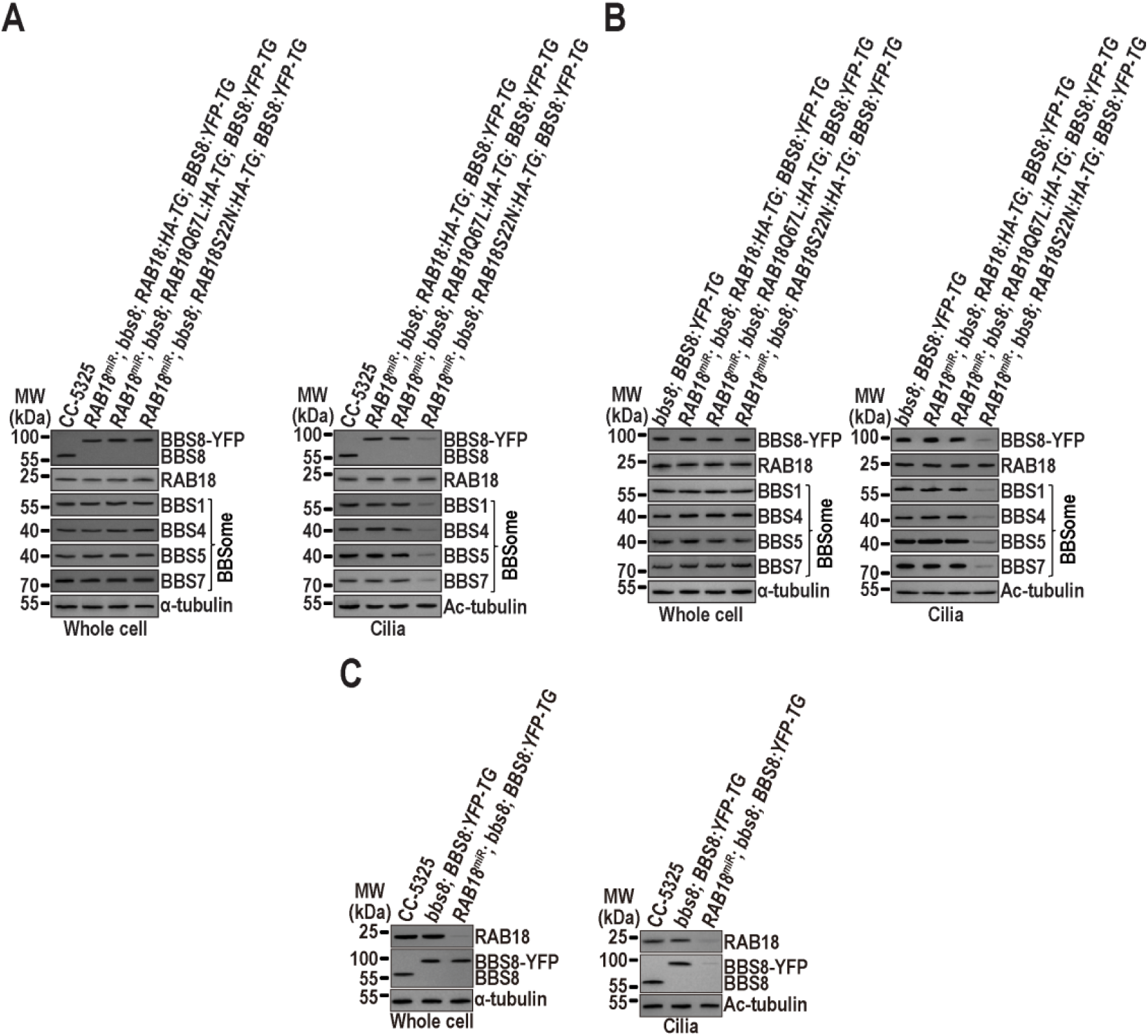
Transgenic strains are screened by immunoblotting. (A). Immunoblots of whole cell samples and cilia from the indicated cells probed with antibodies against BBS8, RAB18, and the BBSome subunits BBS1, BBS4, BBS5, and BBS7. (B). Immunoblots of whole cell samples and cilia from the indicated cells probed with antibodies against BBS8, RAB18, BBS1, BBS4, BBS5, and BBS7. (C). Immunoblots of whole cell samples and cilia from the indicated cells probed with antibodies against RAB18 and BBS8. For all panels, α-tubulin and acetylated (Ac)-tubulin were used to adjust the loading for whole cell samples and cilia, respectively. MW stands for molecular weight.

**Fig. S6.**
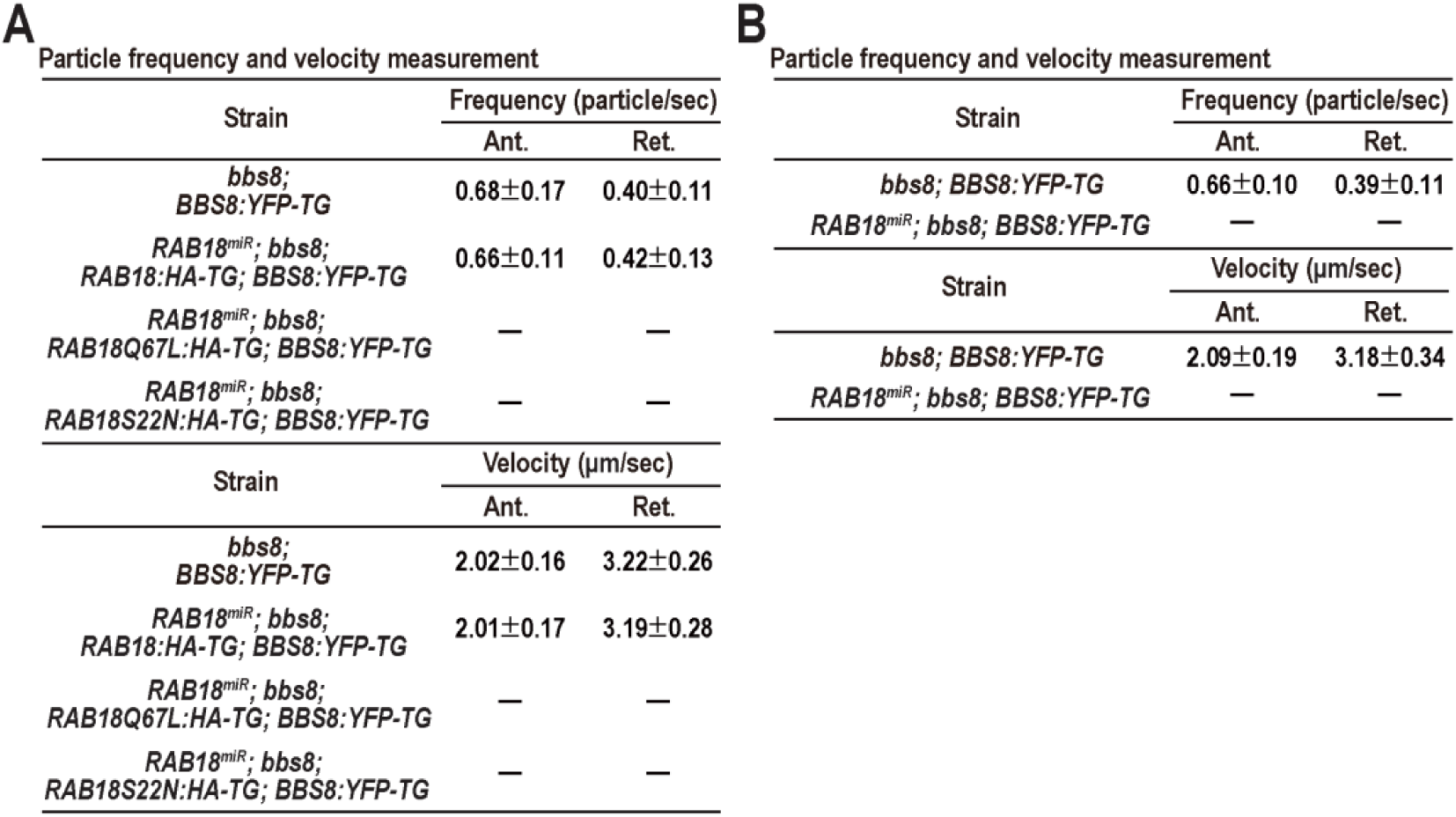
Frequencies and velocities of BBS8-YFP were determined in cilia of various transgenic strains during IFT. (A). Frequencies and velocities of BBS8-YFP in cilia of strains *bbs8; BBS8:YFP-TG*, *RAB18^miR^; bbs8; RAB18:HA-TG; BBS8:YFP-TG*, *RAB18^miR^; bbs8; RAB18Q67L:HA-TG; BBS8:YFP-TG* and *RAB18^miR^; bbs8; RAB18S22N:HA-TG; BBS8:YFP-TG* are presented as numerical values. (B). Frequencies and velocities of BBS8-YFP in cilia of strains *bbs8; BBS8:YFP-TG* and *RAB18^miR^; bbs8; BBS8:YFP-TG* are presented as numerical values. For both tables, “Ant.” and “Ret.” represent anterograde and retrograde, respectively. “-” indicates none of the BBS8-YFP particles observed perform IFT. Data are presented as mean ± S.D. from thirty (frequency) and fifty (velocity) cilia. a one sample unpaired student’s *t*-test is used for statistical analysis.

**Table S1.** Antibodies used in this study.

| Antibody | Dilution |  | Origins | Reference or source |
| --- | --- | --- | --- | --- |
|  | IB | IS |  |  |
| Anti-RAB18 | 1:1,000 | 1:100 | Rb | This study |
| Anti-IFT22 | 1:1,000 | N/A | Rb | (9) |
| Anti-IFT38 | 1:1,000 | N/A | Rb | (7) |
| Anti-IFT43 | 1:500 | N/A | Rb | (12) |
| Anti-IFT46 | 1:1,000 | N/A | Rb | (8) |
| Anti-IFT57 | 1:500 | N/A | Rb | (8) |
| Anti-IFT70 | 1:1,000 | N/A | Rb | (8) |
| Anti-IFT139 | 1:1,000 | N/A | Rb | (8) |
| Anti-BBS1 | 1:1,000 | N/A | Rb | (9) |
| Anti-BBS4 | 1:500 | N/A | Rb | (6) |
| Anti-BBS5 | 1:1,000 | N/A | Rb | (9) |
| Anti-BBS7 | 1:500 | N/A | Rb | (6) |
| Anti-BBS8 | 1:500 | 1:50 | Rb | (7) |
| Anti-PLD | 1:1,000 | 1:25 | Rb | (6) |
| Anti-CEP290 | N/A | 1:50 | RB | (2) |
| Anti-α-tubulin | 1:10,000 | N/A | Mo | Sigma-Aldrich (T5168) |
| Anti-acetylated-tubulin | 1:10,000 | N/A | Mo | Sigma-Aldrich (T7451) |
| Anti-YFP | 1:1,000 | N/A | Mo | Roche (11814460001) |
| Anti-HA | 1:1,000 | N/A | Ra | Roche (11867423001) |
| HRP-conjugated goat anti-rabbit IgG | 1:10,000 | N/A | Gt | The Jackson Lab. (111035003) |
| HRP-conjugated goat anti-mouse IgG | 1:10,000 | N/A | Gt | The Jackson Lab. (115035003) |
| HRP-conjugated goat anti-rat IgG | 1:10,000 | N/A | Gt | The Jackson Lab. (112005003) |
| Alexa-Fluor 488-conjugated goat anti-mouse IgG | N/A | 1:400 | Gt | Molecular Probes (A10680) |
| Alexa-Fluor 594-conjugated goat anti-rabbit IgG | N/A | 1:400 | Gt | Molecular Probes (A11012) |

Note: Rb, rabbit; Mo, mouse; Ra, rat; Gt, goat; HRP, horseradish peroxidase; IB, immunoblotting; IS, immunostaining.

**Table S2.** *Chlamydomonas* strains used in this study.

| Name | Genotype | Reference or source |
| --- | --- | --- |
| CC-5325 | *cw15; mt<sup>-</sup>* | (13) |
| *RAB18; IFT22:HA:YFP-TG* | *nit1; nit2; mt<sup>+</sup>; RAB18:HA:YFP-TG* | (2) |
| *RAB18; IFT38:YFP-TG* | *nit1; nit2; mt<sup>+</sup>; RAB18:HA:YFP-TG* | (2) |
| *RAB18; IFT43:HA:YFP-TG* | *nit1; nit2; mt<sup>+</sup>; RAB18:HA:YFP-TG* | (2) |
| *RAB18<sup>miR</sup>* | *cw15; mt<sup>-</sup>; RAB18<sup>miR</sup>* | This study |
| *RAB18<sup>miR</sup>; IFT22:HA:YFP-TG* | *cw15; mt<sup>-</sup>; RAB18<sup>miR</sup>; IFT22:HA:YFP-TG* | This study |
| *RAB18<sup>miR</sup>; IFT38:YFP-TG* | *cw15; mt<sup>-</sup>; RAB18<sup>miR</sup>; IFT38:YFP-TG* | This study |
| *RAB18<sup>miR</sup>; IFT43:HA:YFP-TG* | *cw15; mt<sup>-</sup>; RAB18<sup>miR</sup>; IFT43:HA:YFP-TG* | This study |
| *RAB18<sup>miR</sup>; RAB18:HA:YFP-TG* | *cw15; mt<sup>-</sup>; RAB18<sup>miR</sup>; RAB18:HA:YFP-TG* | This study |
| *RAB18<sup>miR</sup>; RAB18Q67L:HA:YFP-TG* | *cw15; mt<sup>-</sup>; RAB18<sup>miR</sup>; RAB18Q67L:HA:YFP-TG* | This study |
| *RAB18<sup>miR</sup>; RAB18S22N:HA:YFP-TG* | *cw15; mt<sup>-</sup>; RAB18<sup>miR</sup>; RAB18S22N:HA:YFP-TG* | This study |
| *bbs8* | *bbs8; mt<sup>+</sup>* | (14) |
| *bbs8; BBS8:YFP-TG* | *bbs8; mt<sup>+</sup>; BBS8:YFP-TG* | (2) |
| *RAB18<sup>miR</sup>; bbs8; RAB18:HA:YFP-TG* | *cw15; mt<sup>-</sup>; RAB18<sup>miR</sup>; bbs8; RAB18:HA-TG* | This study |
| *RAB18<sup>miR</sup>; RAB18Q67L:HA:YFP-TG* | *cw15; mt<sup>-</sup>; RAB18<sup>miR</sup>; bbs8; RAB18Q67L:HA-TG* | This study |
| *RAB18<sup>miR</sup>; RAB18S22N:HA:YFP-TG* | *cw15; mt<sup>-</sup>; RAB18<sup>miR</sup>; bbs8; RAB18S22N:HA-TG* | This study |
| *RAB18<sup>miR</sup>; bbs8; RAB18:HA-TG; BBS8:YFP-TG* | *cw15; mt<sup>-</sup>; RAB18<sup>miR</sup>; bbs8; RAB18:HA-TG; BBS8:YFP-TG* | This study |
| *RAB18<sup>miR</sup>; bbs8; RAB18Q67L:HA-TG; BBS8:YFP-TG* | *cw15; mt<sup>-</sup>; RAB18<sup>miR</sup>; bbs8; RAB18Q67L:HA-TG; BBS8:YFP-TG* | This study |
| *RAB18<sup>miR</sup>; bbs8; RAB18S22N:HA-TG; BBS8:YFP-TG* | *cw15; mt<sup>-</sup>; RAB18<sup>miR</sup>; bbs8; RAB18S22N:HA-TG; BBS8:YFP-TG* | This study |
| *HR:YFP-TG* | *nit1; nit2; mt<sup>+</sup>; HA:YFP-TG* | (15) |
| *rab18-876* | *cw15; rab18::aphVIII; mt<sup>-</sup>* | CLiP (13)<br>(LMJ.RY0402.132876) |

**Table S3.**
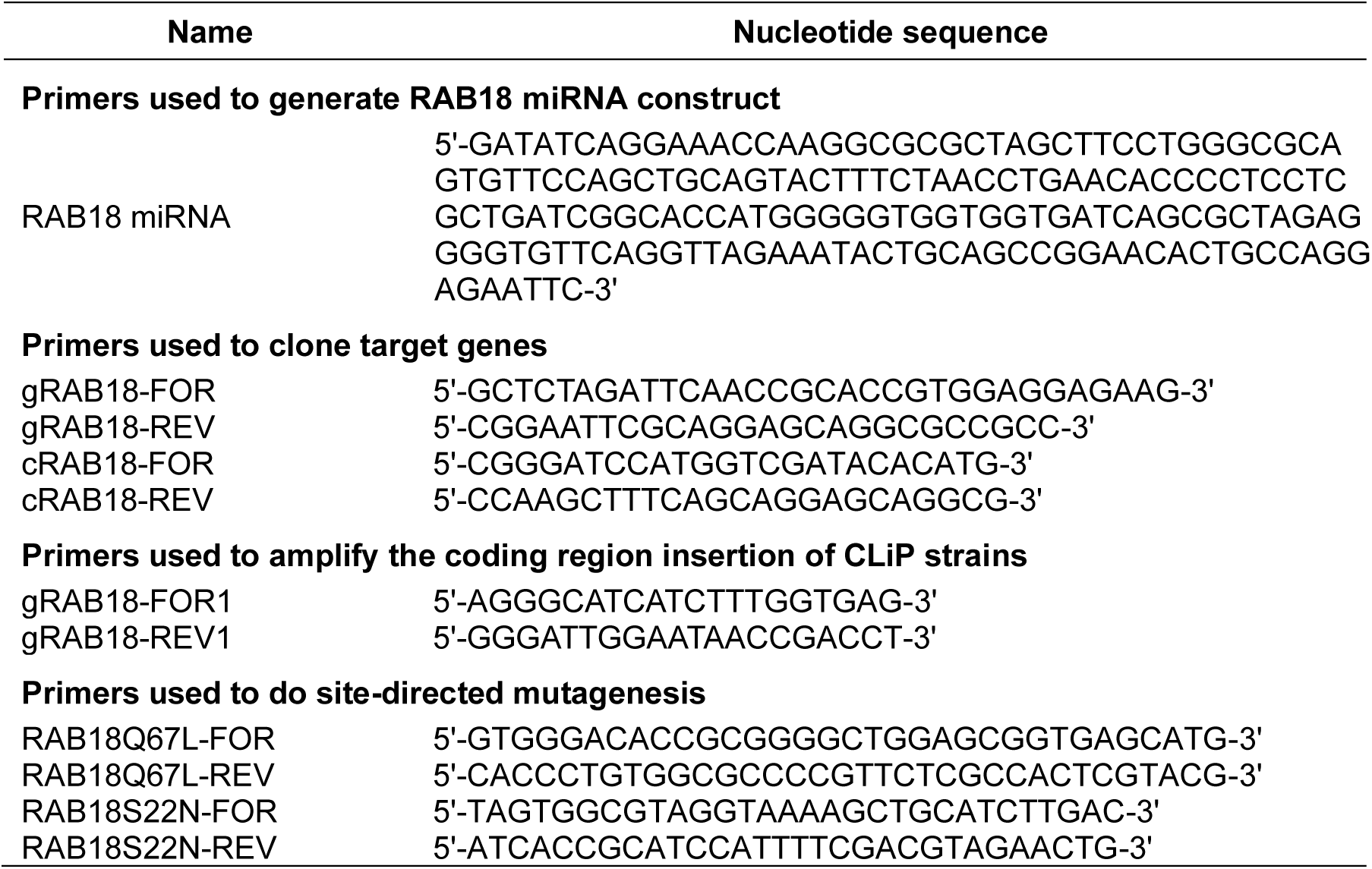
Primers used in this study.

**Movie S1.** TIRF imaging of IFT43-HA-YFP movement in cilia of *RAB18; IFT43:HA:YFP-TG*. A frame from this movie and kymograph are shown in Figure 1C. Play speed is real-time (15 fps).

**Movie S2.** TIRF imaging of IFT43-HA-YFP movement in cilia of *RAB18^miR^; IFT43:HA:YFP-TG*. A frame from this movie and kymograph are shown in Figure 1C. Play speed is real-time (15 fps).

**Movie S3.** TIRF imaging of IFT22-HA-YFP movement in cilia of *RAB18; IFT22:HA:YFP-TG*. A frame from this movie and kymograph are shown in Figure 1C. Play speed is real-time (15 fps).

**Movie S4.** TIRF imaging of IFT22-HA-YFP movement in cilia of *RAB18^miR^; IFT22:HA:YFP-TG*. A frame from this movie and kymograph are shown in Figure 1C. Play speed is real-time (15 fps).

**Movie S5.** TIRF imaging of IFT38-YFP movement in cilia of *RAB18; IFT38:YFP-TG*. A frame from this movie and kymograph are shown in Figure 1C. Play speed is real-time (15 fps).

**Movie S6.** TIRF imaging of IFT38-YFP movement in cilia of *RAB18^miR^; IFT38:YFP-TG*. A frame from this movie and kymograph are shown in Figure 1C. Play speed is real-time (15 fps).

**Movie S7.** TIRF imaging of RAB18-HA-YFP movement in *RAB18^miR^; RAB18:HA:YFP-TG* cilia. A frame from this movie and kymograph are shown in Figure 1F. Play speed is real-time (15 fps).

**Movie S8.** TIRF imaging of RAB18Q67L-HA-YFP movement in cilia of *RAB18^miR^; RAB18Q67L:HA:YFP-TG*. A frame from this movie and kymograph are shown in Figure 2D. Play speed is real-time (15 fps).

**Movie S9.** TIRF imaging of RAB18S22N-HA-YFP movement in cilia of *RAB18^miR^; RAB18S22N:HA:YFP-TG*. A frame from this movie and kymograph are shown in Figure 2D. Play speed is real-time (15 fps).

**Movie S10.** TIRF imaging of BBS8-YFP movement in cilia of *bbs8; BBS8:YFP-TG*. A frame from this movie and kymograph are shown in Figure 3E. Play speed is real-time (15 fps).

**Movie S11.** TIRF imaging of BBS8-YFP movement in the cilia of *RAB18^miR^; bbs8; RAB18:HA-TG; BBS8:YFP-TG*. A frame from this movie and kymograph are shown in Figure 3E. Play speed is real-time (15 fps).

**Movie S12.** TIRF imaging of BBS8-YFP movement in cilia of *RAB18^miR^; bbs8; RAB18Q67L:HA- TG; BBS8:YFP-TG*. A frame from this movie and kymograph are shown in Figure 3E. Play speed is real-time (15 fps).

**Movie S13.** TIRF imaging of BBS8-YFP movement in cilia of *RAB18^miR^; bbs8; RAB18S22N:HA- TG; BBS8:YFP-TG*. A frame from this movie and kymograph are shown in Figure 3E. Play speed is real-time (15 fps).

**Movie S14.** TIRF imaging of RAB18-HA-YFP movement in cilia of *RAB18^miR^; bbs8; RAB18:HA:YFP-TG*. A frame from this movie and kymograph are shown in Figure 5B. Play speed is real-time (15 fps).

**Movie S15.** TIRF imaging of RAB18Q67L-HA-YFP movement in cilia of *RAB18^miR^; bbs8; RAB18Q67L:HA:YFP-TG*. A frame from this movie and kymograph are shown in Figure 5B. Play speed is real-time (15 fps).

**Movie S16.** TIRF imaging of RAB18S22N-HA-YFP movement in cilia of *RAB18^miR^; bbs8; RAB18S22N:HA:YFP-TG*. A frame from this movie and kymograph are shown in Figure 5B. Play speed is real-time (15 fps).

**Movie S17.** TIRF imaging of BBS8-YFP movement in cilia of *bbs8; BBS8:YFP-TG*. A frame from this movie and kymograph are shown in Figure 5D. Play speed is real-time (15 fps).

**Movie S18.** TIRF imaging of BBS8-YFP movement in cilia of *RAB18^miR^; bbs8; BBS8:YFP-TG*. A frame from this movie and kymograph are shown in Figure 5D. Play speed is real-time (15 fps).

## Notes

### Competing Interest Statement

The authors have declared no competing interest.

